# Astrocytes gate long-term potentiation in hippocampal interneurons

**DOI:** 10.1101/2023.06.20.545820

**Authors:** Weida Shen, Yejiao Tang, Jing Yang, Linjing Zhu, Wen Zhou, Liyang Xiang, Feng Zhu, Jingyin Dong, Yicheng Xie, Ling-Hui Zeng

## Abstract

Long-term potentiation is involved in physiological processes such as learning and memory, motor learning and sensory processing, and pathological conditions such as addiction. In contrast to the extensive studies on the mechanism of long-term potentiation on excitatory glutamatergic synapses onto excitatory neurons (LTP_E→E_), the mechanism of LTP on excitatory glutamatergic synapses onto inhibitory neurons (LTP_E→I_) remains largely unknown. In the central nervous system, astrocytes play an important role in regulating synaptic activity and participate in the process of LTP_E→E_, but their functions in LTP_E→I_ remain incompletely defined., We studied the role of astrocytes in regulating LTP_E→I_ in the hippocampal CA1 region and their impact on cognitive function using electrophysiological, pharmacological, confocal calcium imaging, chemogenetics and behavior tests. We showed that LTP_E→I_ in the stratum oriens of hippocampal CA1 is astrocyte independent. However, in the stratum radiatum, synaptically released endocannabinoids increase astrocyte Ca^2+^ via type-1 cannabinoid receptors, stimulate D-serine release, and potentiate excitatory synaptic transmission on inhibitory neurons through the activation of (N-methyl-D-aspartate) NMDA receptors. We also revealed that chemogenetic activation of astrocytes is sufficient for inducing NMDA-dependent *de novo* LTP_E→I_ in the stratum radiatum of the hippocampus. Furthermore, we found that disrupting LTP_E→I_ by knocking down γCaMKII in interneurons of the stratum radiatum resulted in dramatic memory impairment. Our findings suggest that astrocytes release D-serine, which activates NMDA receptors to regulate LTP_E→I_, and that cognitive function is intricately linked with the proper functioning of this LTP_E→I_ pathway.

## Introduction

Long-term potentiation (LTP) was originally identified as a long-term increase in synaptic connections at glutamatergic synapses onto granule cells of the hippocampus (Bliss & Lomo 1973). This form of plasticity has been well documented in CA1 pyramidal cells, where synaptic plasticity can be induced by the activation of postsynaptic (N-methyl-D-aspartate) NMDA receptors (NMDARs), voltage-dependent Ca^2+^ channels, and metabotropic glutamate receptors (mGluRs) (Bliss & Collingridge 1993; Nicoll 2017). Numerous studies have provided compelling evidence that LTP in the hippocampal network occurs not only at excitatory glutamatergic synapses onto excitatory pyramidal and granule cells (LTP_E→E_) but also at excitatory glutamatergic synapses onto inhibitory interneurons (LTP_E→I_) (Kullmann & Lamsa 2007; Kullmann & Lamsa 2011; Asgarihafshejani *et al*. 2022; Le Duigou *et al*. 2015; Pelletier & Lacaille 2008). However, the mechanism underlying LTP_E→I_ remains controversial due to the heterogeneous types of interneurons in the hippocampus and the absence of synaptic spines between excitatory inputs on interneurons (Pelkey *et al*. 2017; Kullmann & Lamsa 2007). Two distinct forms of LTP_E→I_, namely NMDAR-dependent LTP_E→I_ and NMDAR-independent LTP_E→I_, have been observed in hippocampal interneurons (Kullmann & Lamsa 2007; Kullmann & Lamsa 2011).

Astrocytes are the most prevalent glial cell type in the central nervous system (CNS) and play a critical role in regulating the development and function of the nervous system (Escartin *et al*. 2019; Verkhratsky & Nedergaard 2018; Chaboub & Deneen 2013; Perez-Catalan *et al*. 2021). Astrocytes have multipolar branches with numerous microprocesses that allow them to closely associate with blood vessels, neuronal cell bodies and axons, other glial cells and synapses (Bushong *et al*. 2002; Xie *et al*. 2022; Santello *et al*. 2019). Astrocytes also express various ion channels, transporters and neurotransmitter receptors (Ciappelloni *et al*. 2017; Verkhratsky & Steinhäuser 2000; Verkhratsky & Nedergaard 2018). With these membrane proteins, astrocytes can sense neuronal activity and exhibit increases in intracellular Ca^2+^ in reaction to neurotransmitters, and in turn, they release neuroactive chemicals called gliotransmitters that regulate synaptic transmission and plasticity (Araque *et al*. 2014; Bazargani & Attwell 2016; Khakh & McCarthy 2015; Verkhratsky & Nedergaard 2018), but also please see refer to Hamilton and Attwell 2010 (Hamilton & Attwell 2010). Numerous studies have shown that the release of D-serine, a co-agonist of NMDAR, from astrocytes is capable of enabling LTP in cultures, in slices and in vivo (Henneberger *et al*. 2010; Robin *et al*. 2018; Yang *et al*. 2003; Mothet *et al*. 2006; Panatier *et al*. 2006). Given the growing number of studies demonstrating the direct roles that astrocytes play in regulating LTP_E→E_, understanding whether and how astrocytes modulate LTP of excitatory postsynaptic currents in interneurons is of particular interest.

In this particular study, our focus was on the interneurons distributed in the stratum radiatum layer of the CA1 region of the hippocampus. Approximately 80% of these interneurons exhibit NMDAR-dependent LTP_E→I_ when presynaptic stimulation is paired with postsynaptic depolarization. We found that blocking astrocyte metabolism and clamping of astrocyte Ca^2+^ signaling can prevent LTP_E→I_ in large NMDAR-containing interneurons, which could be rescued by bath application of D-serine. Furthermore, pharmacological and Ca^2+^ imaging studies have shown that astrocytes respond to Schaffer collateral stimulation with Ca^2+^ increases through activation of type-1 cannabinoid receptors (CB1Rs), which stimulate the release of D-serine and further regulate LTP_E→I_ via binding to the glycine site of NMDARs. We also found that activating astrocytes with Gq designer receptors exclusively activated by designer drugs (DREADDs) induced a potentiation of excitatory to inhibitory synapses in the stratum radiatum. Additionally, knockdown of γCaMKII hampered LTP_E→I_ in the stratum radiatum of the hippocampus in brain slices and disrupted contextual fear conditioning memory in vivo. Taken together, these results are the first to indicate that astrocytes are an integral component of a form of long-term synaptic plasticity between glutamatergic neurons and GABAergic interneurons, and that memory is also regulated by LTP_E→I_.

## Results

### Astrocytes play a role in the formation of NMDAR-dependent LTP_E→I_ in the CA1 stratum radiatum

To visualize CA1 stratum radiatum interneurons, we delivered an adeno-associated virus serotype 2/9 (AAV2/9) vector encoding EGFP under the control of the interneuronal mDLx promoter (AAV2/9-mDLx-EGFP) to this region (Dimidschstein *et al*. 2016). Within the virally transduced region, EGFP expression was limited to the interneuron, with high penetrance (>98% of the GAD67 cells expressed EGFP) **(Figure 1-figure supplement 1A-C)** and almost complete specificity (>98% EGFP-positive cells were also GAD67 positive) **(Figure 1-figure supplement 1D)**. These immunostaining results indicate that EGFP expression was limited to the interneurons in the CA1 region of the stratum radiatum.

We next examined whether expressing EGFP in interneurons affects their membrane properties and synaptic transmission. Therefore, we performed whole-cell patch recordings of EGFP^+^ interneurons and putative interneurons in the CA1 stratum radiatum of the hippocampus in the control mice in the presence of the GABAA receptor blocker picrotoxin. We found that the excitability and resting membrane potential were not different between these EGFP^+^ interneurons and putative interneurons **(Figure 1-figure supplement 2A-C)**. Moreover, we also found no difference in spontaneous excitatory postsynaptic currents (sEPSCs) or the paired-pulse ratio (PPR) at 50 ms interpulse intervals between EGFP^+^ interneurons and putative interneurons **(Figure 1-figure supplement 2D-H)**. These results indicate that AAV injection and exogenous protein expression in interneurons have no effect on the membrane properties and baseline synaptic transmission of interneurons.

To avoid some necessary ingredient for LTP_E→I_ induction being diluted from the cytoplasm, a perforate patch-clamp was used to record EPSPs from CA1 stratum radiatum interneurons. After a 10 min baseline recording, we delivered theta burst stimulation (TBS) consisting of 100 Hz stimulation with 25 pulses delivered in six trains separated by 20-second intervals. TBS was applied to induce LTP_E→I_ of the excitatory inputs to CA1 interneurons. The interneurons were depolarized to -10 mV using a voltage-clamp model during TBS delivery. Under this condition, we found that in 8 out of 10 cells LTP_E→I_ was induced for at least 45 min **(Figure 1A-C)**. We repatched 6 of these successful LTP_E_ cells and randomly patched 2 EGFP^+^ cells in the stratum radiatum in the whole-cell voltage-clamp model. The results indicated that all cells showed a linear current-voltage (I-V) curve for AMPAR, as well as a significant component of NMDAR-mediated currents **(Figure 1-figure supplement 3)**. In addition, we found that the induction of LTP_E→I_ was completely blocked by the NMDAR blocker D-AP5 **(Figure 1A-C)**. This result confirmed that LTP_E→I_ in CA1 stratum radiatum interneurons is NMDAR dependent (Lamsa *et al*. 2005; Lamsa *et al*. 2007).

**Figure 1.**
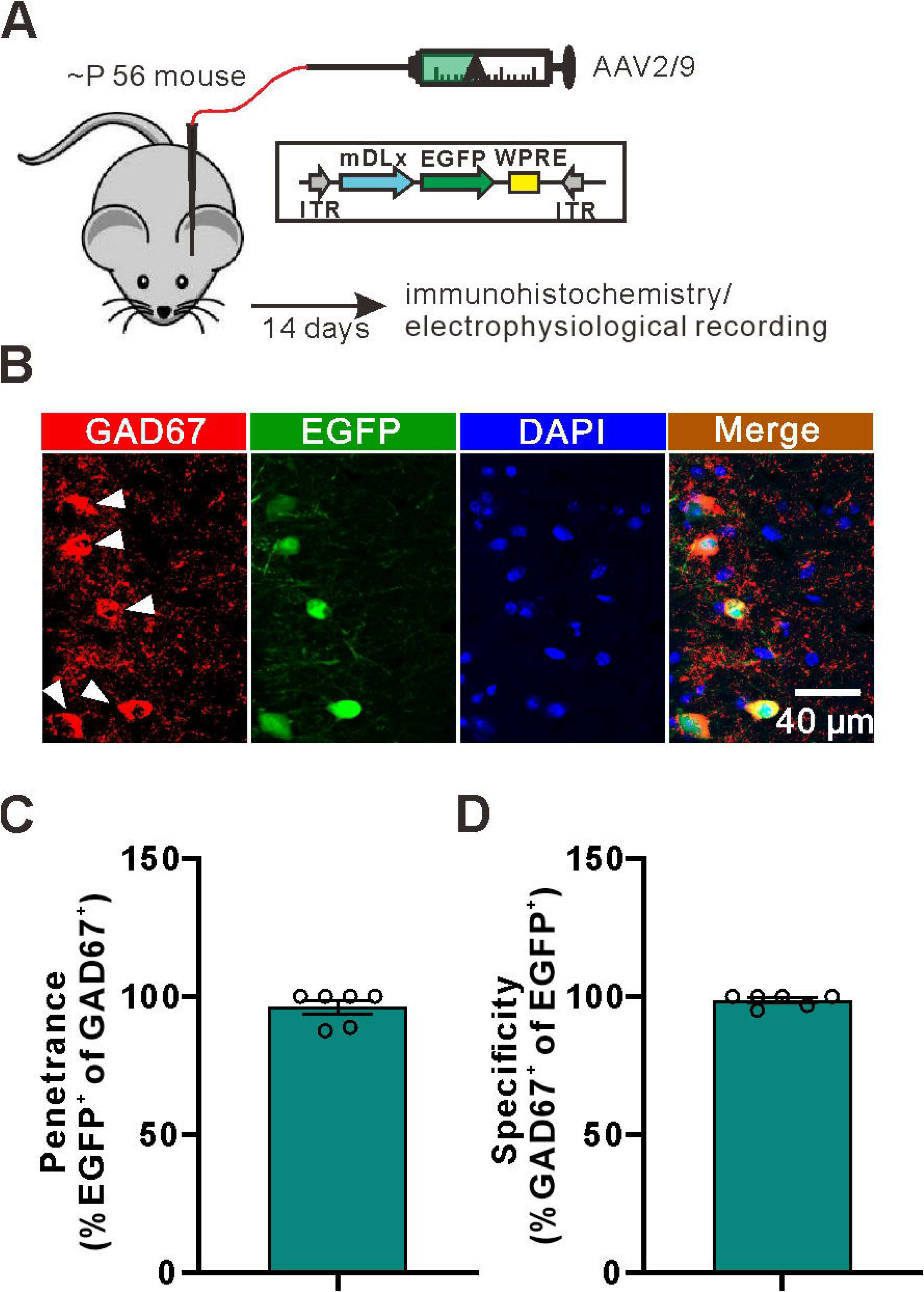
LTP_E→I_ in the stratum radiatum is dependent on the activation of NMDA receptors and astrocytic metabolism. **(A)** Left: superimposed representative averaged EPSPs recorded 10 min before (dark traces) and 35-45 min after (red traces) LTP induction; Right: superimposed representative averaged EPSPs recorded 10 min before (dark traces) and 35-45 min after (red traces) LTP induction in the presence of NMDA receptor antagonist D-AP5. **(B)** Normalized slope before and after the TBS stimulation protocol in control conditions and in the presence of the NMDA receptor antagonist D-AP5. **(C)** The summary data measured 35-45 min after LTP induction **(Control: 220.2±33.94, n=10 slices from 5 mice; D-AP5: 105.2±4.404, n=6 slices from 3 mice; t=2.580 with 14 degrees of freedom, *p*=0.0218, two-tailed unpaired t-test).** **(D)** Left: superimposed representative averaged EPSPs recorded 10 min before (dark traces) and 35-45 min after (red traces) LTP induction; Right: superimposed representative averaged EPSPs recorded 10 min before (dark traces) and 35-45 min after (red traces) LTP induction while astrocytic metabolism was disrupted. **(E)** Normalized slope before and after the TBS stimulation protocol in control conditions and in the presence of the astrocytic metabolism inhibitor FAC. **(F)** The summary data measured 35-45 min after LTP induction **(Control: 193.5±21.84, n=11 slices from 5 mice; FAC: 103.8±7.278, n=6 slices from 6 mice; t=2.943 with 15 degrees of freedom, *p*=0.0101, two-tailed unpaired t-test).**

To investigate whether astrocytes were involved in NMDAR-dependent LTP_E→I_ in CA1 stratum radiatum interneurons, we treated slices with fluoroacetate (FAC, 5 mM) to specifically block astrocyte metabolism (Swanson & Graham 1994; Henneberger *et al*. 2010). The results showed that the induction of LTP_E→I_ was blocked by FAC but not in the control slice **(Figure 1D-F)**. Next, we tested whether glial metabolism is involved in LTP_E→I_ in CA1 stratum orient, which has been shown to depend on calcium-permeable AMPARs (CP-AMPARs) and metabotropic glutamate receptors (mGluRs) (Le Duigou *et al*. 2015). We found that the induction of LTP_E→I_ is not blocked by FAC in the CA1 stratum oriens (**Figure 1-figure supplement 4A-B**). We repatched 6 of these cells (4 cells from the control group and 2 cells from the FAC-treated group) in whole-cell voltage-clamp mode and observed that they exhibited high rectification AMPARs and a small component of NMDAR-mediated current (**Figure 1-figure supplement 5**). This observation confirms findings from a previous study (Lamsa *et al*. 2007; Oren *et al*. 2009). Moreover, we found that the induction of LTP_E→I_ is not blocked by D-AP5 in the CA1 stratum oriens (**Figure 1-figure supplement 4**). Overall, the results indicate that the formation of LTP_E→I_ in the CA1 stratum radiatum is tightly regulated by astrocyte function. The results in the stratum oriens also exclude the possibility that FAC directly affects the metabolism of interneurons which inhibit the formation of LTP_E→I_.

It is commonly accepted that astrocytic calcium signaling plays a pivotal role in triggering the release of gliotransmitters and modulating synaptic transmission (Bazargani & Attwell 2016; De Pitta *et al*. 2016; Sancho *et al*. 2021; Navarrete *et al*. 2012; Goenaga *et al*. 2023). In addition, it is well documented that astrocytic calcium signaling is necessary for the formation of LTP_E→E_ in the CA1 stratum radiatum (Henneberger *et al*. 2010; Robin *et al*. 2018). Thus, we tested whether intracellular Ca^2+^ signals are needed for the induction of LTP_E_ in the CA1 stratum radiatum. We found that clamping of the astrocyte Ca^2+^ concentration significantly suppressed LTP_E→I_ **(Figure 2A-D)**. Consistent with the previous study on LTP_E→E_, the supply of D-serine (50 μM) fully rescued NMDAR-dependent LTP_E→I_ **(Figure 2D)**. For the control, when the intracellular Ca^2+^ concentration of astrocytes was not clamped but the astrocytes were recorded with a glass pipette, LTP_E→I_ was indistinguishable from that induced without patching an astrocyte **(Figure 2D)**. Overall, these results indicate that functional preservation of astrocytic metabolism and Ca^2+^ mobilization is critical for maintaining the induction of LTP_E→I_.

**Figure 2.**
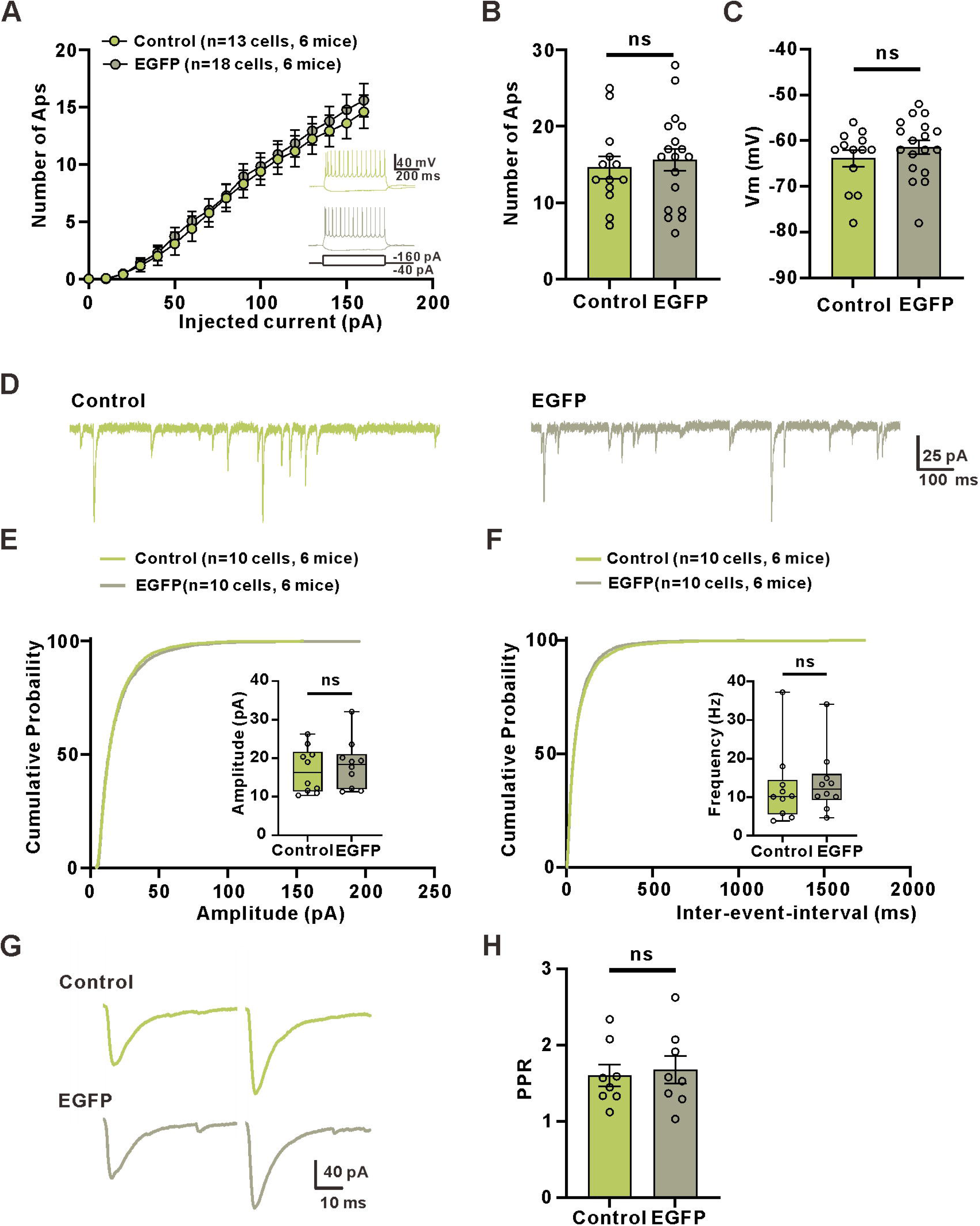
Astrocyte Ca^2+^ is involved in the induction of LTP_E→I_. **(A)** Representative image of the astrocytic syncytium illustrating that Ca^2+^ clamping reagents and Alexa Fluor 594 diffuse through gap junctions surrounding the patched interneuron. **(B)** Left: superimposed representative averaged EPSPs recorded 10 min before (dark traces) and 35-45 min after (red traces) LTP induction; Right: superimposed representative averaged EPSPs recorded 10 min before (dark traces) and 35-45 min after (red traces) LTP induction when the Ca^2+^ concentration was clamped. **(C)** Normalized slope before and after the TBS stimulation protocol in control conditions and when Ca^2+^ concentration was clamped. **(D)** The summary data measured 35-45 min after LTP induction **(ACSF: 162.7±22.21, n=6 slices from 3 mice; Ca^2+^ clamp: 111.1±7.323, n=7 slices from 4 mice; Mann-Whitney U Statistic= 3.000, *p*=0.008, Mann-Whitney Rank Sum Test; ACSF: 197.5±33.83, n=6 slices from 4 mice; Ca^2+^ clamp+D-serine: 195.1±31.75, n=7 slices from 4 mice; Mann-Whitney U Statistic= 19.000, *p*=0.836, Mann-Whitney Rank Sum Test; ACSF: 172.2±19.49, n=8 slices from 4 mice; patch: 189.9±38.11, n=6 slices from 3 mice; Mann-Whitney U Statistic= 22.000, *p*=0.852, Mann-Whitney Rank Sum Test)**.

### Astrocytic Ca^2+^ transients, induced by activation of astroglial cannabinoid type 1 receptors (CB1Rs), are involved in the regulation of LTP_E_**_→_**_I_ formation

Accumulating evidence indicates that neuronal depolarization in the hippocampus induces astrocytic Ca^2+^ transients, which are mediated by the activation of astroglial CB1Rs (Eraso-Pichot *et al*. 2023; Noriega-Prieto *et al*. 2023; Navarrete & Araque 2008; Navarrete & Araque 2010; Navarrete *et al*. 2014). Moreover, astroglial CB1R-mediated Ca^2+^ elevation is necessary for LTP_E→E_ in the CA1 region of the hippocampus (Robin *et al*. 2018). Therefore, we asked whether astroglial CB1R-mediated Ca^2+^ elevations are needed for hippocampal LTP_E→I_. We first analyzed whether TBS could evoke astrocytic Ca^2+^ transients via CB1Rs in the stratum radiatum of the hippocampus. In this study, GCaMP6f was used to analyze astrocytic Ca^2+^ signals, which were specifically expressed in astrocytes by using adeno-associated viruses of the 2/5 serotype (AAV 2/5) with the astrocyte-specific gfaABC_1_D promoter **(Figure 3-figure supplement 1A)**. Furthermore, the expression was confirmed by immunohistochemistry. Within the virally transduced region, GCaMP6f-positive cells in the CA1 stratum radiatum were also positive for the astrocyte-specific marker GFAP **(Figure 3-figure supplement 1A and C-D)**. Costaining with the neuron marker NeuN showed no overlap with GCaMP6f expression **(Figure 3-figure supplement 1B, E)**. Consistent with previous studies (Sherwood *et al*. 2017; Robin *et al*. 2018), we found that TBS significantly increased Ca^2+^ signaling in astrocytes of hippocampal slices in the presence of picrotoxin and CGP55845 **(Figure 3A-B and F)**. As predicted, the increase in Ca^2+^ signals after TBS was inhibited by the CB1 receptor inhibitor AM251 (2 μM) **(Figure 3C-D and F)**. Previous studies have demonstrated that activation of [1-adrenoceptors increases calcium signals (Shen *et al*. 2021; Ding *et al*. 2013; Gordon *et al*. 2005; Bekar *et al*. 2008; Paukert *et al*. 2014; Oe *et al*. 2020) and triggers the release of D-serine release in the neocortex (Pankratov & Lalo 2015). However, our results show that [1-adrenoceptor receptors are not involved in the Ca^2+^ signal increase observed after TBS **(Figure 3F)**.

**Figure 3.**
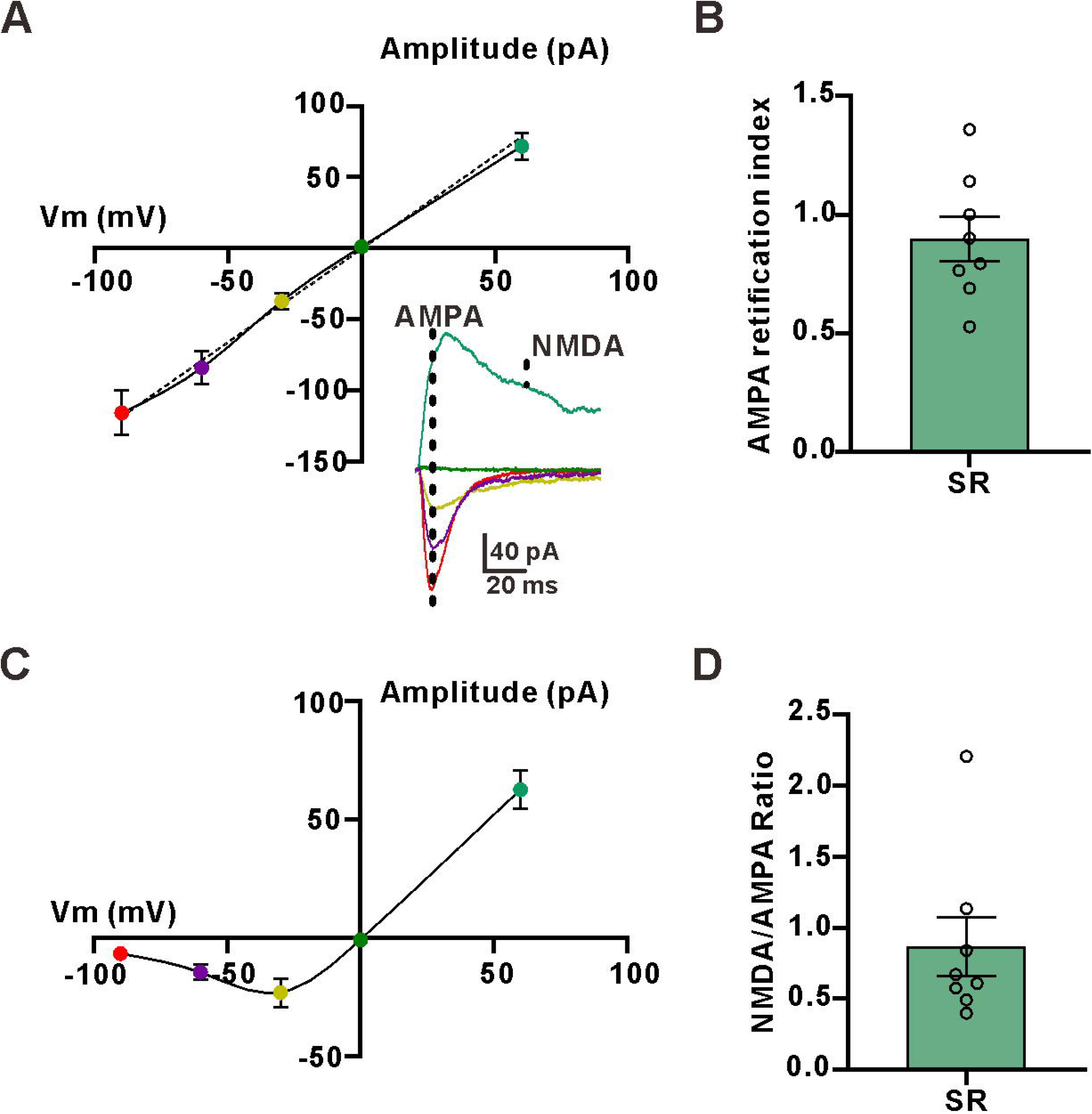
Activation of astrocytic CB1 receptors causes an increase in astrocytic Ca^2+^ signals and is involved in the induction of LTP_E→I_. **(A)** Representative images of a GCaMP6f^+^ astrocyte before (left) and after (right) theta burst stimulation (TBS). **(B)** Kymographs and _△_F/F traces of cells with Ca^2+^ signals evoked by the activation of Schaffer collateral with TBS in GCaMP6f^+^ astrocytes. **(C)** Representative images of a GCaMP6f^+^ astrocyte before (left) and after (right) theta burst stimulation (TBS) in the presence of the CB1 receptor blocker AM251(2 μM). **(D)** Kymographs and _△_F/F traces of cells with Ca^2+^ signals evoked by the activation of Schaffer collateral with TBS in GCaMP6f^+^ astrocytes in the presence of the CB1 receptor blocker AM251. **(E)** Kymographs and _△_F/F traces of cells with Ca^2+^ signals evoked by the activation of Schaffer collateral with a TBS in GCaMP6f^+^ astrocytes in the presence of the α1-adrenoceptor blocker terazosin. **(F)** Summary plots for experiments in (B)-(E) illustrating that Ca^2+^ signals elicited by TBS are mediated by the activation of CB1 receptors **(Control pre: 0.3275±0.1083** _△_ F/F**, Control post: 1.313±0.2483** _△_F/F**, n=45 ROI from 20 cells of 5 mice; z= 3.369, *p*<0.001, Wilcoxon Signed Rank Test; AM251 pre: 0.3541±0.07219** _△_F/F**, AM251 post: 0.3418±0.07240** _△_ F/F**, n=28 ROI from 13 cells from 5 mice; z= -0,797, *p*=0.432, Wilcoxon Signed Rank Test; Terazosin pre: 0.5725±0.1749** _△_ F/F**, Terazosin post: 1.73±0.4661** _△_F/F**, n=25 ROI from 11 cells from 4 mice; z=3.215, *p*=0.001, Wilcoxon Signed Rank Test).** **(G)** Upon: superimposed representative averaged EPSPs recorded 10 min before (dark traces) and 35-45 min after (red traces) LTP induction; below: superimposed representative averaged EPSPs recorded 10 min before (dark traces) and 35-45 min after (red traces) LTP induction in the presence of the CB1 receptor antagonist AM251. **(H)** Normalized slope before and after the TBS stimulation protocol in control conditions and in the presence of the CB1 receptor antagonist AM251. **(I)** The summary data measured 35-45 min after LTP induction **(Control: 180.2±13.76, n=12 slices from 5 mice; AM251: 118.9±7.406, n=10 slices from 4 mice; t=3.697 with 20 degrees of freedom, *p*=0.0014, two-tailed unpaired t-test; Control: 169.3±16.89, n=12 slices from 6 mice; AM251+D-serine: 177.4±14.27, n=10 slices from 5 mice; t=0.3561 with 20 degrees of freedom, *p*=0.7255, two-tailed unpaired t-test).**

Next, we explored whether CB1R-mediated Ca^2+^ elevation was accompanied by the formation of LTP_E→I_. We found that LTP_E→I_ was significantly reduced in AM251-treated slices relative to that in controls **(Figure 3G-I)**. Consistent with the above results shown in **Figure 2D**, LTP_E→I_ was rescued by the addition of D-serine in AM251-treated slices

### D-serine release from astrocytes potentiates the NMDAR-mediated synaptic response

Next, we explored the underlying mechanisms by which astrocytes control the formation of LTP_E→I_ in the CA1 stratum radiatum. Our above results indicate that D-serine is a downstream signaling pathway of astrocyte Ca^2+^ signaling. D-serine, a co-agonist of NMDAR, can be released by astrocytes through Ca^2+^-dependent exocytosis and regulate the function of NMDAR. It has been shown that the occupancy of synaptic NMDAR co-agonist sites by D-serine is not saturated in CA1 pyramidal cells (Robin *et al*. 2018; Papouin *et al*. 2012). It has been shown that the NMDAR co-agonist site in CA1 pyramidal neurons is fully saturated during the dark phase, but this saturation dissipates to subsaturating levels during the light phase (Papouin *et al*. 2017a). However, the level of occupancy of synaptic NMDAR co-agonist sites by D-serine in interneurons of the CA1 stratum radiatum remains unclear. Furthermore, previous studies have demonstrated that NMDAR-dependent synaptic responses of interneurons in the stratum radiatum display a strong rundown effect under whole-cell mode (Lamsa *et al*. 2005). Thus, we recorded the NMDAR-mediated EPSPs from stratum radiatum interneurons in the perforated-patch configuration in Mg^2+^-free ACSF, which showed no rundown effect in this configuration **(Figure 4A-B)**. Bath application of 50 μM D-serine enhanced the NMDAR-mediated EPSPs **(Figure 4A-C)**, indicating that the level of D-serine in the excitatory to inhibitory synapse cleft was not saturated to occupy the co-agonist site. Subsequently, we employed a similar protocol in another study to test whether TBS could increase the extracellular level of D-serine and promote NMDAR-mediated synaptic responses (Henneberger *et al*. 2010). Our results indicated that NMDAR-mediated responses were transiently enhanced after one TBS, and the percentage of potentiation was reduced by pretreatment of hippocampal slices with 50 μM D-serine **(Figure 4D-F)**. However, disrupting glial metabolism with FAC rendered the percentage of TBS-induced potentiation not significantly different for that in the group pretreated with 50 μM D-serine **(Figure 4F)**. Furthermore, the percentage of potentiation in AMPAR-mediated responses after TBS was insensitive to bath application of D-serine, indicating that the enhancement of the NMDAR-mediated response by TBS was not due tochanges in release probability and cell excitability **(Figure 4F)**. Overall, our results indicate that D-serine release from astrocytes potentiate NMDAR-mediated responses by binding the co-agonist site and regulating the formation of LTP_E→I_.

**Figure 4.**
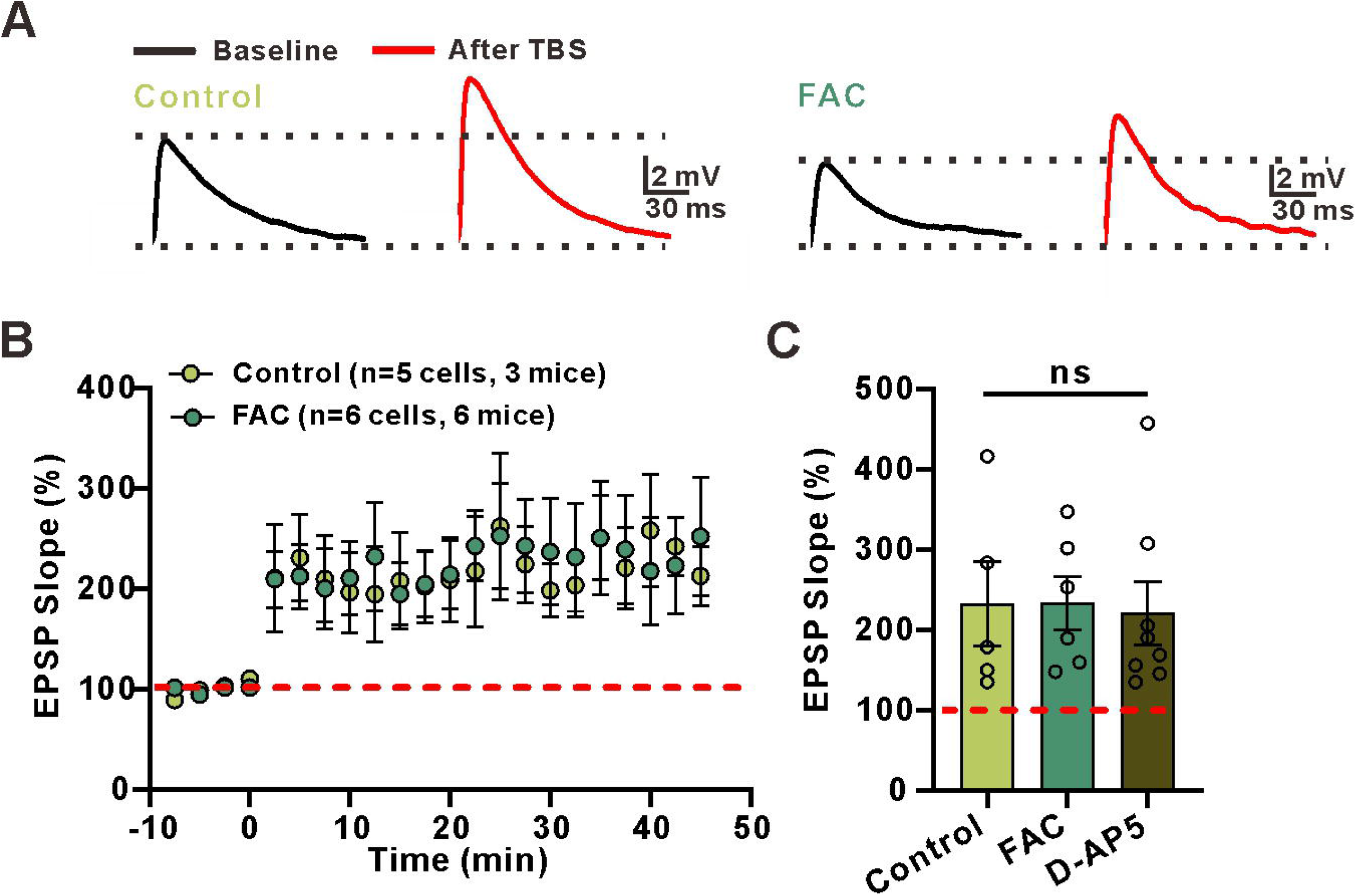
D-Serine release from astrocytes potentiates NMDAR-mediated responses. **(A)** Example traces of NMDAR-mediated EPSPs before and after bath application of D-serine in stratum radiatum interneurons. **(B)** Normalized amplitude before and after addition of D-serine in stratum radiatum interneurons **(C)** The percentage of potentiation in NMDAR-mediated responses **(percentage of potention: 155.523 ± 11.234, n=6 slices from 3 mice; t=-4.942 with 6 degrees of freedom, p=0.00260, two-tailed paired t-test).** **(D)** Left: example traces of NMDAR-mediated EPSPs before and after one episode of the TBS stimulation protocol in stratum radiatum interneurons. Right: example traces of NMDAR-mediated EPSPs before and after the TBS stimulation protocol in stratum radiatum interneuronsin the presence of D-serine. **(E)** Normalized amplitude before and after the TBS stimulation protocol in stratum radiatum interneurons during ASCF and after application of D-serine. **(F)** The summary data measured the percentage of potentiation in NMDAR-mediated responses **(Control: 155.5±11.23, n=7 slices from 3 mice; D-serine: 121.1±6.917, n=8 slices from 4 mice; FAC: 112.8±4.931, n=7 slices from 3 mice; *p*=0.0036, F(2, 19)=7.653, ANOVA with Dunnett’s comparison) and AMPAR-mediated responses after TBS(Control: 132.357±12.305, n=5 slices from 3 mice; D-serine: 136.012±10.611, n=8 slices from 4 mice; t=-0.224 with 10 degrees of freedom, p=0.827, two-tailed unpaired t-test).**

### Chemogenetic activation of astrocytes induced E**_→_**I synaptic potentiation

The crucial role of astrocytes in LTP_E→E_ has been extensively demonstrated in brain slices (Henneberger et al., 2010; Min and Nevian, 2012; Pascual et al., 2005; Perea and Araque, 2007; Suzuki et al., 2011) and in vivo (Robin *et al*. 2018). Previous studies have demonstrated that the activation of astrocytes through chemogenetics leads to potentiation at CA1 synapses in the hippocampus in the absence of high-frequency stimulation (Nam *et al*. 2019; Adamsky *et al*. 2018; Van Den Herrewegen *et al*. 2021). In this study, we aimed to investigate whether the activation of the astrocytic G protein-coupled receptor (GPCR) pathway could trigger the potentiation of E→I synaptic transmission. We utilized an adeno-associated virus serotype 5 (AAV2/5) vector encoding hM3Dq fused to mCherry for specific activation of astrocytes via clozapine-N-oxide (CNO, 5 μM). To ensure specific expression in astrocytes, the vector was also under the control of the astrocyte-specific gfaABC1D promoter **(Figure 5-figure supplement 1)**.

To verify whether CNO application could evoke Ca^2+^ transients in astrocytes, we delivered AVV of hM3Dq along with AAV of GCaMP6f and conducted confocal Ca^2+^ imaging in brain slices. Our results showed that CNO application indeed induced an increase in intracellular Ca^2+^ levels in cells coexpressing hM3Dq and GCaMP6f **(Figure 5A-D)**. These results suggest that the expression of hM3Dq is selective to astrocytes and can elicit a rise in intracellular Ca^2+^ levels upon administration of CNO. Subsequently, we investigated the impact of astrocytic Gq activation on evoked synaptic events in interneurons of the CA1 stratum radiatum that were induced by Schaffer collaterals stimulation, both before and after the administration of CNO. Interestingly, we observed that the EPSC amplitude was potentiated by 60% in response to the exact same stimulus in gfaABC1D::hM3Dq slices treated with CNO **(Figure 5E-G)**, while no such potentiation was detected in slices obtained from the mice that were injected with a control virus (AAV2/5-gfaABC1D::mCherry) **(Figure 5E-G**).

**Figure 5.**
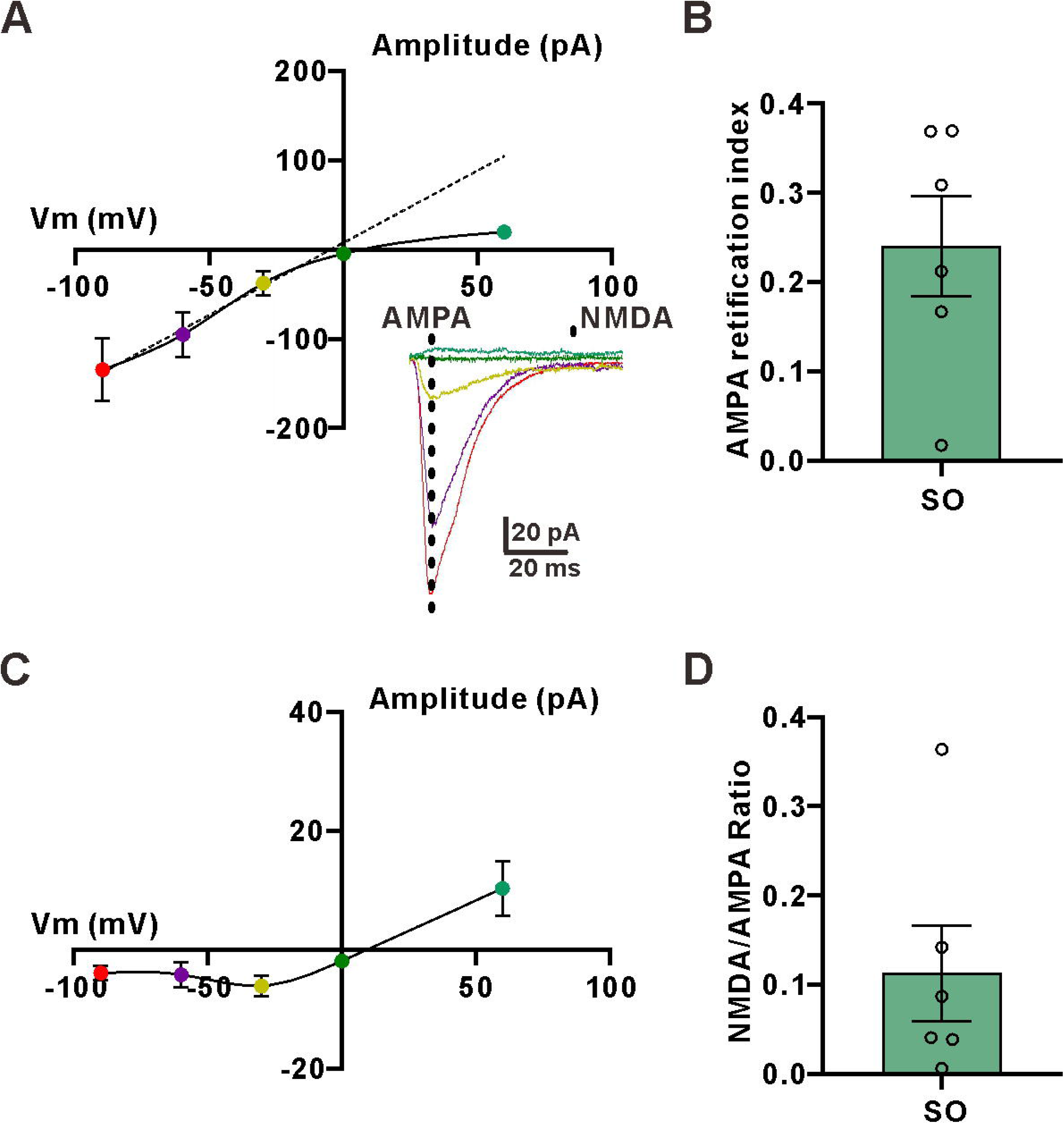
Astrocytic activation induces *de novo* LTP_E→I_. **(A)** Representative images of a GCaMP6f^+^ astrocyte before (left) and after (right) CNO (5 μM) application. **(B)** Representative confocal images showing coexpression of GCaMP6f and mCherry constructs confirm that the representative cell in (A) coexpressed hM3Dq. **(C)** Kymographs and _△_ F/F traces of cells with Ca^2+^ signals evoked by bath application of CNO in GCaMP6f^+^ and mCherry^+^ astrocytes. **(D)** Summary plot illustrating that CNO is effective in activating astrocytes **(Baseline: 0.6167±0.07762** _△_F/F**, CNO: 1.456±0.1504** _△_F/F**, n=36 from 19 cells of 3 mice; t=6.411 with 35 degrees of freedom, *p*<0.0001, two-tailed paired t-test)**. **(E)** Upon: superimposed representative averaged EPSPs recorded 10 min before (dark traces) and 35-45 min after (red traces) CNO application when astrocytes only express mCherry; below: superimposed representative averaged EPSPs recorded 10 min before (dark traces) and 35-45 min after (red traces) CNO application when hM3Dq-expressed astrocytes were activated by CNO. **(F)** Relative EPSP slope before and after CNO application when astrocytes only express mCherry and when astrocytes express hM3Dq. **(G)** The summary data measured 35-45 min after LTP induction **(mCherry: 100.6±5.651, n=5 slices from 3 mice; hM3Dq: 172.8±10.28, n=7 slices from 4 mice; t=5.474 with 10 degrees of freedom, *p*=0.0003, two-tailed unpaired t-test).** **(H)** Upon: superimposed representative averaged EPSPs recorded 10 min before (dark traces) and 35-40 min after (red traces) CNO application; middle: superimposed representative averaged EPSPs recorded 10 min before (dark traces) and 35-40 min after (red traces) CNO application when in the presence NMDA receptor blocker D-AP5; below: superimposed representative averaged EPSPs recorded 10 min before (dark traces) and 35-40 min after (red traces) CNO application when the glycine sites of NMDA receptors were saturated by D-serine. **(I)** Relative EPSP slope before and after CNO application when astrocytes were activated by CNO and in the presence of the NMDA receptor antagonist D-AP5 and in the presence of D-serine. **(J)** The summary data measured 35-45 min after LTP induction **(hM3Dq+CNO: 197.1±27.57, n=6 slices from 3 mice; hM3Dq+CNO+D-AP5: 101.3±6.886, n=6 slices from 3 mice; hM3Dq+CNO+D-serine: 104.2+8.166, n=7 slices from 4 mice; *p*=0.0011, F(2, 16)=10.80, ANOVA with Dunnett’s comparison).**

Previous studies have indicated that the synaptic potentiation triggered by chemogenetic activation of astrocytes is mediated through the release of D-serine by astrocytes, resulting in the activation of NMDARs (Adamsky *et al*. 2018). To verify whether the astrocytic-induced synaptic potentiation between excitatory and inhibitory neurons is mediated by D-serine release from astrocytes and the subsequent activation of NMDARs, we conducted an experiment in which we administered CNO after blocking the NMDARs with D-AP5 or saturating glycine site of NMDARs with 50 μM D-serine. Our results showed that both the NMDAR blocker D-AP5 and 50 μM D-serine completely inhibited the potentiation in EPSP amplitude observed in response to CNO-induced astrocytic activation **(Figure 5H-J)**. Our findings demonstrate, for the first time, that astrocytic activation alone can trigger *de novo* potentiation of synapses between excitatory and inhibitory neurons and that this potentiation is indeed mediated by the release of D-serine from astrocytes and subsequent activation of NMDARs.

### LTP_E_**_→_**_I_ in CA1 of the stratum radiatum is necessary for long-term memory formation

We have previously shown that GABAergic interneurons in the hippocampus express high levels of γCaMKII, while αCaMK[, βCaMK [and δCaMK [are expressed at a lower frequency (He *et al*. 2021). Additionally, studies have demonstrated that γCaMKII expressed in hippocampal parvalbumin-positive (PV^+^) interneurons and cultured hippocampal inhibitory interneurons is essential for the induction of LTP_E→I_ (He *et al*. 2021; He *et al*. 2022). Specifically, in hippocampal PV^+^ interneurons, this protein is also vital for the formation of hippocampus-dependent long-term memory in vivo (He *et al*. 2021). Therefore, we asked whether astrocyte-gated LTP_E→I_ in the CA1 stratum radiatum is involved in the hippocampus-dependent long-term memory formation. Several independent groups have provided compelling evidence suggesting that astroglial CB1R-mediated signaling pathways regulate excitatory synapses formed by excitatory neurons rather than excitatory synapses between excitatory neurons and interneurons (Fernandez-Moncada & Marsicano 2023; Eraso-Pichot *et al*. 2023; Noriega-Prieto *et al*. 2023; Kano *et al*. 2009; Navarrete & Araque 2008; Navarrete & Araque 2010; Han *et al*. 2012). Thus, manipulating this pathway may also affect excitatory synapses formed by excitatory neurons. To avoid this side effect, we specifically knocked downγCaMKII expression in inhibitory neurons of the CA1 stratum radiatum by delivering an adeno-associated virus serotype 2/9 (AAV2/9) vector encoding shRNAs against γCaMKII and EGFP under the control of an inhibitory neuron-specific promoter (AAV2/9-mDLx-EGFP-γCaMKII shRNA) through bilateral stereotactic injection. We observed that interneurons identified by the specific marker GAD67 were also positive for EYFP and therefore likely expressed shRNAs, which led to knockdown γCaMKII in these cells **(Figure 6-figure supplement 1)**. To confirm that knockdown of γCaMKII could hamper the induction of LTP_E→I_, EYFP-positive interneurons in the CA1 stratum radiatum were recorded in perforated-patch mode. We found that knocking down γCaMKII in CA1 stratum radiatum interneurons impaired LTP_E→I_, but LTP_E→I_ in the putative interneurons in the stratum radiatum of control mice was unimpaired **(Figure 6A-C)**. In addition, robust LTP_E→I_ was induced by TBS in EYFP-positive interneurons injected with AAV-mDLx-scramble shRNA into the CA1 stratum radiatum of the hippocampus **(Figure 6C)**. Moreover, it is worth noting that the resting membrane potential, frequency and amplitude of sEPSCs, excitability and PPR recorded from γCaMKII knockdown interneurons did not differ from those recorded from putative interneurons and EYFP-positive interneurons (infected with scramble shRNA) **(Figure 6-figure supplement 2)**. Taken together, these results indicate that knocking down γCaMKII from interneurons has no effect on synaptic transmission at baseline but impairs the induction of LTP_E→I_ in these interneurons.

**Figure 6.**
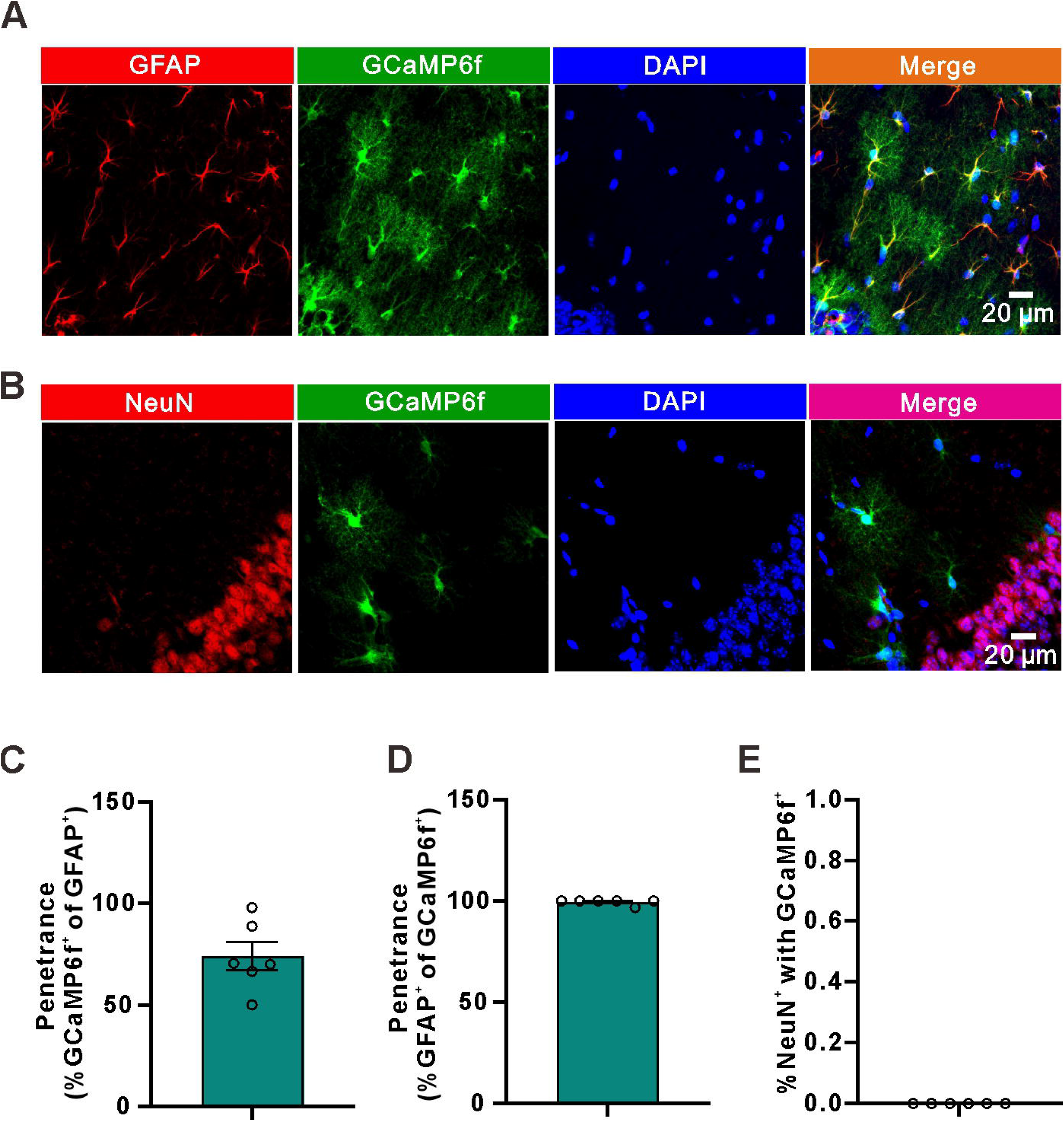
Impaired hippocampus-dependent long-term memory in γCaMKII knockdown mice. **(A)** Upon: superimposed representative averaged EPSPs recorded 10 min before (dark traces) and 35-45 min after (red traces) LTP induction in the WT group; middle: superimposed representative averaged EPSPs recorded 10 min before (dark traces) and 35-45 min after (red traces) LTP induction in the AAV-mDLx-scramble shRNA group; below: superimposed representative averaged EPSPs recorded 10 min before (dark traces) and 35-45 min after (red traces) LTP induction in the AAV-mDLx-γCaMKII shRNA group. **(B)** Normalized slope before and after the TBS stimulation protocol in the control, scramble shRNA and γCaMKII shRNA groups. **(C)** The summary data measured 35-45 min after LTP induction **(Control: 196±21.56, n=12 slices from 5 mice;** γ**CaMKII shRNA: 121.3±10.64, n=14 slices from 6 mice; scramble shRNA: 199.6±21.42, n=11 slices from 5 mice; *p*=0.0040, F(2, 34)=6.527, ANOVA with Dunnett’s comparison).** **(D)** Schematic illustration of the experimental design for contextual and cued fear conditioning test, indicating the timeline of the experimental manipulations. **(E)** The freezing responses was measured in context A before training in WT, bilaterally injected γCaMKII shRNA and scramble shRNA mice **(Control: 17.5±2.975, n=10 mice;** γ**CaMKII shRNA: 20.9±2.275, n=15 mice; scramble shRNA: 20.29±2.341, n=15 mice; *p*=0.6378, F(2, 37)=0.4553, ANOVA with Dunnett’s comparison).** **(F)** The freezing responses were measured 24 h after training in WT, bilaterally injected γCaMKII shRNA and scramble shRNA mice **(Control: 52.4±6.695, n=10 mice;** γ**CaMKII shRNA: 30.64±4.574, n=15 mice; scramble shRNA: 49.27±4.899, n=15 mice; *p*=0.0105, F(2, 37)=5.170, ANOVA with Dunnett’s comparison).** **(G)** The freezing response were measured 28 h after training in WT, bilaterally injected γCaMKII shRNA and scramble shRNA mice **(Pre Control: 19.5±2.171, n=10 mice; Pre** γ**CaMKII shRNA: 25.01±3.016, n=15 mice; Pre scramble shRNA: 21.94±3.519, n=15 mice; *p*=0.4986, F(2, 37)=0.7093, ANOVA with Dunnett’s comparison; Tone 1 Control: 57.9±5.153, n=10 mice; Tone 1** γ**CaMKII shRNA: 56.09±3.319, n=15 mice; Tone 1 scramble shRNA: 55.32±3.368, n=15 mice; *p*=0.9002, F(2, 37)=0.1055, ANOVA with Dunnett’s comparison; Tone 2 Control: 59.75±3.825, n=10 mice; Tone 2** γ**CaMKII shRNA: 60.95±3.482, n=15 mice; Tone 2 scramble shRNA: 60.34±4.485, n=15 mice; *p*=0.9802, F(2, 37)=0.01996, ANOVA with Dunnett’s comparison).**

Next, we examined the behavioral consequences of destroying astrocyte-gated LTP_E→I_ in the CA1 stratum radiatum. We analyzed contextual fear conditioning memory, which is associated with activation of the hippocampus. We found that no effect of γCaMKII knockdown on exploration of the context before conditioning was observed **(Figure 6D-E)**. 24 hours after training, the mice were returned to the training box, and freezing was measured during the first 2 min. We found that γCaMKII knockdown mice showed a significant reduction in freezing during contextual conditioning **(Figure 6F)**. Consistent with our previous study (He *et al*. 2021), γCaMKII knockdown in interneurons in the CA1 stratum radiatum did not produce a significant effect on freezing during tone conditioning at 28 h after training **(Figure 6G)**. Taken together, our results strongly suggest that the astroglial CB1R signaling pathway gated LTP_E→I_ in the CA1 stratum radiatum plays a vital role in hippocampus-dependent long-term memory.

## Discussion

In the present study, we found that LTP_E→I_ in the CA1 stratum radium is tightly controlled by D-serine release from astrocytes via the astroglial CB1R-mediated Ca^2+^ elevation. In addition, knockdown of γCaMKII by specific shRNA in interneurons in the CA1 stratum radium causes cognitive function deficits. Taken together, our data indicate that astrocyte-gated LTP_E→I_ in the CA1 stratum radium plays a critical role in preserving normal cognitive function.

CB1Rs are widely expressed in various brain regions, including the hippocampus, and are detectable in presynaptic terminals (Castillo *et al*. 2012; Kano *et al*. 2009), postsynaptic terminals (Marinelli *et al*. 2009; Bacci *et al*. 2004), intracellular organelles (Jimenez-Blasco *et al*. 2020; Gutierrez-Rodriguez *et al*. 2018) and astrocytes (Ramon-Duaso *et al*. 2023; Noriega-Prieto *et al*. 2023; Eraso-Pichot *et al*. 2023; Fernandez-Moncada & Marsicano 2023; Robin *et al*. 2018; Han *et al*. 2012; Navarrete & Araque 2010; Navarrete & Araque 2008). However, the effects of astroglial CB1R-mediated signaling on synaptic transmission and plasticity are highly debated. It has been shown that exogenous administration of Δ^9^-tetrahydro-cannabinol (THC) leads to temporally prolonged and spatially widespread activation of astroglial CB1 receptors and triggers glutamate release, which activates postsynaptic NMDARs and induces LTD in the CA3–CA1 hippocampal synapses, resulting in working memory deficits (Han *et al*. 2012). Araque and colleagues reported that eCB released from depolarized CA1 pyramidal cells activates astroglial CB1Rs with in a shorter and localized manner and induces glutamate release, which activates lateral presynaptic mGluRs and induces LTP (Navarrete & Araque 2010; Navarrete & Araque 2008). In another study, Robin et al. found that high-frequency stimulation (HFS) of Schaffer Collateral induces LTP, which is gated by the activation of astroglial CB1Rs and the release of D-serine from astrocytes (Robin *et al*. 2018). These diverse consequences of CB1 receptor activation may be due to different neuronal activity patterns that induce eCB release and the different nature of the agonists. Interestingly, it has been discovered that individual hippocampal astrocytes are capable of releasing both ATP/adenosine and glutamate and that this release occurs in a time-dependent and activity-sensitive manner in response to neuronal interneuron activity (Covelo & Araque 2018). These findings suggest that the specific type and intensity of astrocyte stimulation plays a critical role in determining the downstream signaling pathways that are triggered by CB1R activation in astrocytes. Consistent with a previous study, we found that the activation of astroglial CB1Rs induces Ca^2+^ elevation and triggers the release of D-serine which binds to postsynaptic NMDARs and induces LTP formation (Robin *et al*. 2018). Our results provide evidence that astroglial CB1R-mediated signaling not only modulates the E→E synapses, but also regulates E→I synapses.

The precise mechanisms by which neurons and astrocytes differentially regulate D-serine levels are yet to be fully elucidated (Papouin *et al*. 2017b; Wolosker *et al*. 2017; Wolosker *et al*. 2016). However, astrocytes play a significant role in regulating the availability of D-serine. The enzyme serine racemase catalyzes the conversion of L-serine into D-serine, which was initially found in astrocytes and microglia in the mammalian brain (Panatier *et al*. 2006; Stevens *et al*. 2003; Wolosker *et al*. 1999). It should be highlighted that serine racemase has also been detected in neurons (Benneyworth *et al*. 2012; Miya *et al*. 2008; Dun *et al*. 2008). A study showed that, despite a significant reduction in SR protein levels in the brains of neuronal SR knockout mouse brains, the reduction in D-serine levels was minimal, suggesting that neurons are not the exclusive source of D-serine (Benneyworth *et al*. 2012) and that neurons may produce and release D-serine under certain conditions. Notably, activation of G protein-coupled receptors in astrocytes through chemogenetic methods leads to LTP, which relies on the release of D-serine from astrocytes and the activation of NMDARs (Van Den Herrewegen *et al*. 2021; Adamsky *et al*. 2018). Specifically, the release of D-serine, which is regulated by CB1R in astrocytes, is needed for Ca^2+^-dependent modulation of LTP in vivo (Robin *et al*. 2018), as well as the threshold and amplitude of dendritic spikes (Bohmbach *et al*. 2022). Moreover, recent studies have shown that conditional connexin double knockout (Hosli *et al*. 2022) or knockdown of α4nAChR (Ma *et al*. 2022) in astrocytes can decrease the extracellular concentration of D-serine, which in turn reduces NMDAR-dependent synaptic potentiation. These findings suggest that astrocytes are the primary source of D-serine, which plays a crucial role in modulating the function of NMDARs.

It is well established that LTP_E→E_ observed in the CA1 region of the hippocampus is triggered by activation of NMDARs. However, LTP_E→I_ is less studied than LTP_E→E_, but recent evidence suggests that it also involves the activation of NMDARs (He *et al*. 2021; Lamsa *et al*. 2007; Kullmann & Lamsa 2007; Kullmann & Lamsa 2011; Lamsa *et al*. 2005; Nissen *et al*. 2010). There have been reports of NMDAR-dependent LTP_E→I_ in various regions of the brain, including the hippocampus and cortex (Kullmann & Lamsa 2007; Kullmann & Lamsa 2011). Our results suggest that different synaptic mechanisms are involved in the induction of LTP_E→I_ in different subregions of the hippocampus. The stratum radiatum, where interneurons contain NMDARs, is known to be sensitive to NMDAR-dependent LTP_E→I_. In contrast, the stratum oriens, where interneurons contain CP-AMPARs, appears to rely on the activation of CP-AMPA receptors for LTP_E→I_ induction. Notably, our findings suggest that astrocytes contribute to NMDAR signaling in the induction of LTP_E→I_ in the stratum radiatum through the release of the co-agonist D-serine. Notably, our study found that prolonged activation of astrocytes via the Gq-DREADD pathway resulted in a substantial and persistent increase in Ca^2+^ events and significantly potentiated EPSP responses. This is consistent with earlier observations made by other groups regarding LTP_E→E_ (Adamsky *et al*. 2018; Van Den Herrewegen *et al*. 2021). Above all, the mechanism of LTP_E→I_ in the stratum radiatum appears to be shared by LTP_E→E_ observed in the CA1 region.

A previous study demonstrated that knocking down γCaMKII from interneurons can disrupt LTP_E→I_ and cognitive function (He *et al*. 2021; He *et al*. 2022). Our results confirmed that knocking down γCaMKII in interneurons of the stratum radiatum also leads to disruption of LTP_E→I_ and cognitive function. Ma and colleagues showed that, following learning, hippocampal network oscillations in the gamma and theta bands were significantly weaker in γCaMKII knockout mice than in wild-type mice(He *et al*. 2021). This finding suggests that impaired experience-dependent oscillations in the hippocampus of γCaMKII PV-KO mice may lead to cognitive dysfunction. In this respect, it will be intriguing to investigate the network oscillation after learning in our condition in future studies.

In the hippocampus, GABAergic local circuit inhibitory interneurons make up approximately 10-15% of the total neuronal cell population (Bezaire & Soltesz 2013). However, these interneurons are diverse in their subtypes, morphology, distribution, and functions (Pelkey *et al*. 2017; Booker & Vida 2018). In our study, we mainly focused on a subpopulation of interneurons in the stratum radiatum of the hippocampus. Although it is unclear which type of interneuron was recorded in our study, our study indicated that most interneurons in the stratum radiatum do not express CP-AMPARs but express an abundance of NMDARs. These findings are in line with a previous study conducted by Lasmsa et al. (Lamsa *et al*. 2007). In our study, we observed that approximately 80% of interneurons in the stratum radium were able to induce LTP successfully. This finding contrasts with the observation made by Lamsa et al., who reported a figure of approximately 52% interneurons capable of inducing LTP. The reason for this discrepancy could be attributed to differences in the induction protocol used in the respective studies.

Our study corroborates earlier research that suggests that distinct synaptic mechanisms are involved in LTP induction in the CA1 region of the hippocampus across different subregions (Le Duigou *et al*. 2015; Lamsa *et al*. 2007; Kullmann & Lamsa 2011). However, the major breakthrough of our study is the demonstration that astrocytic function serves as the gating mechanism for LTP _E→I_ induction in the stratum radiatum. Additionally, our data reveal that the activation of astrocytes via the Gq-DREADD pathway produces *de novo* long-lasting potentiation of EPSP in stratum radiatum interneurons and that the knockdown of γCaMKII disrupts cognitive function. These results shed light on the complex mechanisms underlying learning and memory in the hippocampus and may have implications for developing new therapies targeted at modulating astrocytic function for the treatment of memory disorders.

## Materials and Methods

### Animals

Our study was conducted in accordance with the Guide for the Care and Use of Laboratory Animals and was approved by the ethics committee of Hangzhou City University (registration number: 22061). C57BL/6 male mice (2-4 months) were purchased from Hangzhou Ziyuan Laboratory Animal Corporation and housed in groups of three to four per cage. The mice were maintained on a 12-hour light/dark cycle and were provided with *ad libitum* access to food and water.

### Stereotactic virus injection

Stereotactic virus injection was conducted as described previously (Shen *et al*. 2021; Shen *et al*. 2022). Briefly, adult mice were deeply anesthetized with sodium pentobarbital (50 mg/kg) and secured in a stereotaxic device with ear bars (RWD, 68930), while their body temperature was maintained at approximately 37 °C using a heating blanket. Their hair was removed using a razor, and the skin was sterilized with iodophor. A 1-cm incision in the midline was made using sterile scissors. Small burr holes were drilled bilaterally using an electric hand drill at the following coordinates: anteroposterior (AP), 2.3 mm from bregma; mediolateral (ML), ±1.4 mm. Virus particles were then injected bilaterally into the stratum radiatum (1.2 mm from the pial surface) using glass pipettes connected to an injection pump (RWD, R480). The injection rate was controlled at 1 nl/s using the pump. To allow the virus to disseminate into the tissue, a glass pipette was left in place for 10 minutes after each injection. After the injection, the pipettes were gradually removed, and the wound was sutured. For hippocampal interneuron physiological recording, 200 nl AAV2/9 mDLx EGFP (6.4 × 10^11^ gc/ml) was injected. For hippocampal slice Ca^2+^ imaging, 500 nl AAV2/5 GfaABC_1_D GCaMP6f (1.2 × 10^12^ gc/ml) was injected alone or mixed with 500 nl AAV2/5 GfaABC_1_D hM3D (Gq) mCherry (5.3 × 10^12^ gc/ml). To knockdown γCaMKII in hippocampal interneurons, a shRNA sequence (5’-GCAGCTTGCATCGCCTATATC-3’) was used. A total of 200 nl of AAV2/9 mDLx γCaMKII shRNA (6.4 × 10^12^ gc/ml) was injected. All viruses were generated by Brainvta and Sunbio Medical Biotechnology (Wuhan, https://www.brainvta.tech/ and Shanghai, http://www.sbo-bio.com.cn/). Two to three weeks after viral injection, the mice were utilized for subsequent experiments.

### Electrophysiology

Mice were anesthetized with isoflurane, and their brains were quickly extracted and immersed in an ice-cold solution which containing (in mM) 235 sucrose, 1.25 NaH_2_PO_4_, 2.5 KCl, 0.5 CaCl_2_, 7 MgCl_2_, 20 glucose, 26 NaHCO_3_, and 5 pyruvate (pH 7.3, 310 mOsm, saturated with 95% O_2_ and 5% CO_2_) at 10-11:00 (UTC[+[08:00) in the morning. Coronal hippocampal slices (300-350 μm) were prepared with a vibrating slicer (Leica, V T1200) and incubated for 30-40 min at 32 °C in artificial cerebrospinal fluid (ACSF) containing (in mM) 26 NaHCO_3_, 2.5 KCl, 126 NaCl, 20 D-glucose, 1 sodium pyruvate, 1.25 NaH_2_PO_4_, 2 CaCl_2_ and 1 MgCl_2_ (pH 7.4, 310 mOsm, saturated with 95% O_2_ and 5% CO_2_).

The slices were transferred to an immersed chamber and continuously perfused with oxygen-saturated ACSF and GABA receptor blockers, picrotoxin (100 μM) and CGP55845 (5 µM), at a rate of 3 ml/min. Interneurons in the dorsal hippocampus of the stratum radiatum or stratum oriens were visualized using infrared differential interference contrast and epifluorescence imaging. Perforated-path recordings were conducted as previously described (Liu *et al*. 2017). Briefly, perforated whole-cell recordings from stratum radiatum or stratum oriens interneurons were made with pipettes filled with solution containing (in mM) 136 K-gluconate, 9 NaCl, 17.5 KCl, 1 MgCl_2_, 10 HEPES, 0.2 EGTA, 25 μM Alexa 488, amphotericin B (0.5 mg ml^−1^) and small amounts of glass beads (5–15 μm in diameter; Polysciences, Inc., Warminster, PA, USA) (pH 7.3, 290 mOsm). The patched neuron was intermittently imaged with epifluorescence to monitor dye penetration. If the patch ruptured spontaneously, the experiment was discontinued.

Whole-cell voltage-clamp recordings were made from either stratum radiatum or stratum oriens interneurons using pipettes with resistance of 3-4 MΩ and filled with a solution containing (in mM) 4 ATP-Na_2_, 0.4 GTP-Na, 125 CsMeSO_3_, 10 EGTA, 10 HEPES, 5 4-AP, 8 TEA-Cl, 1 MgCl_2_, 1 CaCl_2_ (pH 7.3–7.4, 280–290 mOsm).

Whole-cell current-clamp recordings were made from stratum interneurons using pipettes with a resistance of 4-6 MΩ and filled with a solution containing (in mM) 125 K-Gluconate, 2 MgCl_2_, 10 HEPES, 0.4 Na^+^-GTP, 4 ATP-Na_2_, 10 Phosphocreatine disodium salt, 10 KCl, 0.5 EGTA (pH 7.3–7.4, 280–290 mOsm).

Whole-cell recordings were performed on stratum radiatum astrocytes using pipettes with a resistance of 8-10 MΩ and filled with an intracellular solution containing (in mM) 130 K-Gluconate, 20 HEPES, 3 ATP-Na_2_, 10 D-Glucose, 1 MgCl_2_, 0.2 EGTA (pH 7.3-7.4, 280-290 mOsm). In a subset of experiments, 0.14 mM CaCl_2_ and 0.45 mM EGTA were included in the upper intracellular solution to maintain a stable level of astrocytic concentration (calculation by Web-MaxChelator) (Shen *et al*. 2022; Henneberger *et al*. 2010). Astrocytes were identified as described previously (Shen *et al*. 2021; Shen *et al*. 2022).

Electrical stimuli were delivered via theta glass pipettes in the Schaffer Collateral of the stratum radiatum, with a 30 s intertrial interval during baseline (10 min) and after LTP induction (45 min). Evoked EPSPs were recorded in the current-clamp model at the resting membrane potential of the stratum radiatum interneuron. After a 10-min stable baseline period, LTP was induced by applying theta-burst stimulation [TBS; five bursts at 200-ms intervals (5 Hz), each burst consisting of five pulses at 100 Hz]. Each burst was paired with 60-ms long depolarizing steps to -10 mV. Six episodes of TBS paired with depolarization were given at 20-s intervals. In the stratum oriens, LTP was induced by applying TBS paired with five 60-ms long hyperpolarizing steps to -90 mV.

sEPSCs were recorded in a whole-cell model at -70 mV in the presence of picrotoxin (100 μM) and D-AP5 (50 μM). Paired pulses were delivered at an interpulse intervals of 50 ms, and the paired-pulse ratio was calculated by dividing the peak amplitude of the second EPSC by the peak amplitude of the first EPSC. To analyze action potential properties, interneurons were recorded at rest and depolarized with 500-ms current injection pulses at 10-pA increments.

To avoid NMDAR-mediated signaling rapidly washing out when recording interneurons in whole-cell mode, electrical stimuli were delivered with a 5 s intertrial interval to measure the I-V relationship for NMDAR-mediated EPSPs within 5 minutes of breaking in. The NMDA/AMPA ratio in the stratum radiatum or stratum oriens interneurons was calculated by measuring the amplitude of NMDAR-mediated EPSCs at +60 mV (50 ms after stimulation) and the peak amplitude of AMPAR-mediated EPSCs recorded at −60 mV. To measure pure NMDAR-mediated EPSCs in interneurons at resting membrane potential in a perforated whole-cell recording model, Mg^2+^-free ACSF was used, and D-AP5 (50 μM) and picrotoxin (100 μM) were present in ACSF.

Axopatch 700B amplifiers were utilized for patch-clamp recordings (Molecular Devices). The data were filtered at 6 kHz and sampled at 20 kHz before being processed off-line using the pClampfit 10.6 program (Molecular Devices). Bridge balances were automatically compensated for in whole-cell current clamp recordings. Series resistance was not compensated, but negative pulses (-10 mV) were employed to monitor series resistances and membrane resistances. The data were included in the analysis if the series resistances varied by less than 20% over the course of the trial. All experiments were carried out at 32 °C.

### Behavioral assays

For the contextual and cued fear conditioning test, the mouse was habituated to the environment and handled for three consecutive days. On the fourth day, the mice were allowed to explore the conditioning cage for 2 min, after which they received three moderate tone-shock pairs [30 stones (80 dB, 4 kHz) coterminating with a foot shock (0.4 mA, 2 s)]. Following conditioning, the mice were returned to their home cages. The next day, they were placed in the same conditioning cage but without receiving any foot shocks, and their freezing behavior was analyzed for the first 5 min. Four hours after the contextual fear conditioning test, the mice were placed in a novel cage with a different shape and texture of the floors compared to those of the conditioning cage. They were allowed to freely explore the new environment for 2 min and then subjected to 3-tone stimulations [(80 dB, 4 kHz) lasting for 30 s] separated by intervals of 90 s, but without receiving any foot-shocks. Once the final tone had ended, all of the mice were allowed to freely explore the chamber for an additional 90 s. Fear responses were measured by calculating the freezing values of the mice using Packwin software (Panlab, Harvard Apparatus, USA). All apparatuses were carefully cleaned with 30% ethanol in between tests.

### Immunohistochemistry

Immunohistochemistry was conducted as described previously (Shen *et al*. 2017; Nikolic *et al*. 2018; Shen *et al*. 2021). Briefly, mice were administered a single intraperitoneal injection of 50 mg/kg sodium pentobarbital for anesthesia and were transcardially perfused with phosphate-buffered saline (PBS). After the liver and lungs had become bloodless, the mice were perfused with 4% paraformaldehyde (PFA) in 0.1 M PBS. The brains were quickly removed and placed in 4% PFA at 4 [overnight. Next, the tissues were cryoprotected in successive concentrations of 10%, 20%, and 30% sucrose before being sliced into 20 μm sections using a freezing microtome. After being washed multiple times with PBS, the sections were blocked for 1.5 hours at room temperature (22-24 [) in a blocking solution consisting of 5% bovine serum albumin (BSA) and 1% Triton X-100. Following the blocking step, the sections were incubated with primary antibodies overnight at 4 [. The following antibodies were used: mouse monoclonal anti-GFAP (1/1000, Cell Signaling Technology Cat #3670, RRID: AB_561049), mouse monoclonal anti-NeuN (1/500, Millipore Cat# MAB377, RRID: AB_2298772), and mouse monoclonal anti-GAD67 (1/500, Synaptic Systems Cat# 198 006, RRID: AB_2713980). After washing several times in PBS, the sections were then incubated with the following secondary antibody: Alexa Fluor 594 goat anti-mouse (1/1000, Cell Signaling Technology Cat# 8890, RRID: AB_2714182) at room temperature for 2 h. Afterward, the sections were rinsed several times in PBS and incubated with DAPI for 5 minutes at room temperature. Next, the sections were rinsed again and mounted with Vectashield mounting medium. Images were examined using a confocal laser scanning microscope (Olympus, VT1000) and analyzed using ImageJ (NIH, RRID: SCR_003070).

### Ca^2+^ imaging

Ca^2+^ signals in hippocampal astrocytes were observed under a confocal microscope (Fluoview 1000; Olympus) with a 40x water immersion objective lens (NA = 0.8). GCaMP6f was excited at 470 nm and the emission signals were further filtered through a 490-582 nm bandpass filter. mCherry was excited at 594 nm and the emission signals were further filtered through a 580-737 nm bandpass filter. Astrocytes located in the hippocampal CA1 region and at least 40 µm away from the slice surface were selected for imaging. Images were acquired at 1 frame per 629 ms. Hippocampal slices were maintained in ACSF containing picrotoxin (100 μM) and CGP55845 (5 µM) using a perfusion system. One episode of TBS was used to stimulate the neuron. CNO (5 μM) was bath-applied to activate hM3D(Gq)-mediated Ca^2+^ signals. The Ca^2+^ signal analysis was described previously (Nikolic *et al*. 2018; Shen *et al*. 2021; Shen *et al*. 2022; Shen *et al*. 2017). In some experiments, image stacks comprising 10-15 optical sections with 1 µM z-spacing were acquired to facilitate the identification of GCaMP6f and hM3Dq (mCherry) coexpression in astrocytes. Briefly, movies were registered using the StackReg plugin of ImageJ to eliminate any x-y drift. Ca^2+^ signals were then analyzed in selected ROIs using the Time Series Analyzer V3 plugin of ImageJ. GCaMP6f fluorescence was calculated as ΔF/F = (F-F_0_)/F_0_. The mean ΔF/F of Ca^2+^ signals wasanalyzed using Clampfit 10.6.

### Statistics

All data processing, figure generation, layout, and statistical analysis were performed using Clampfit 10.6, Prism, MATALAB and Coreldraw. To assess the normality of the data, the Shapiro-Wilk test was used. If the results of the test were not statistically significant (p> 0.05), the data were assumed to follow a normal distribution, and a paired t-test was employed. On the other hand, if the test was statistically significant, a Wilcoxon Signed Rank Test or Mann-Whitney Ran Sum Test was utilized. To statistically analyze cumulative frequency distributions, the Kolmogorov-Smirnov test was employed. When comparing two groups, either a Wilcoxon signed-rank test or a Student’s t-test (paired or unpaired) was used. When comparing three groups, One-way repeated measures (RM) ANOVA followed by Dunnett’s post-hoc test was used.

All values are presented as the mean ± standard error of the mean (SEM). Data were considered significantly different when the p value was less than 0.05 (*p < 0.05, **p < 0.01, ***p < 0.001). The numbers of cells or mice used in each analysis are indicated in the figure legends.

## Key Resources Table

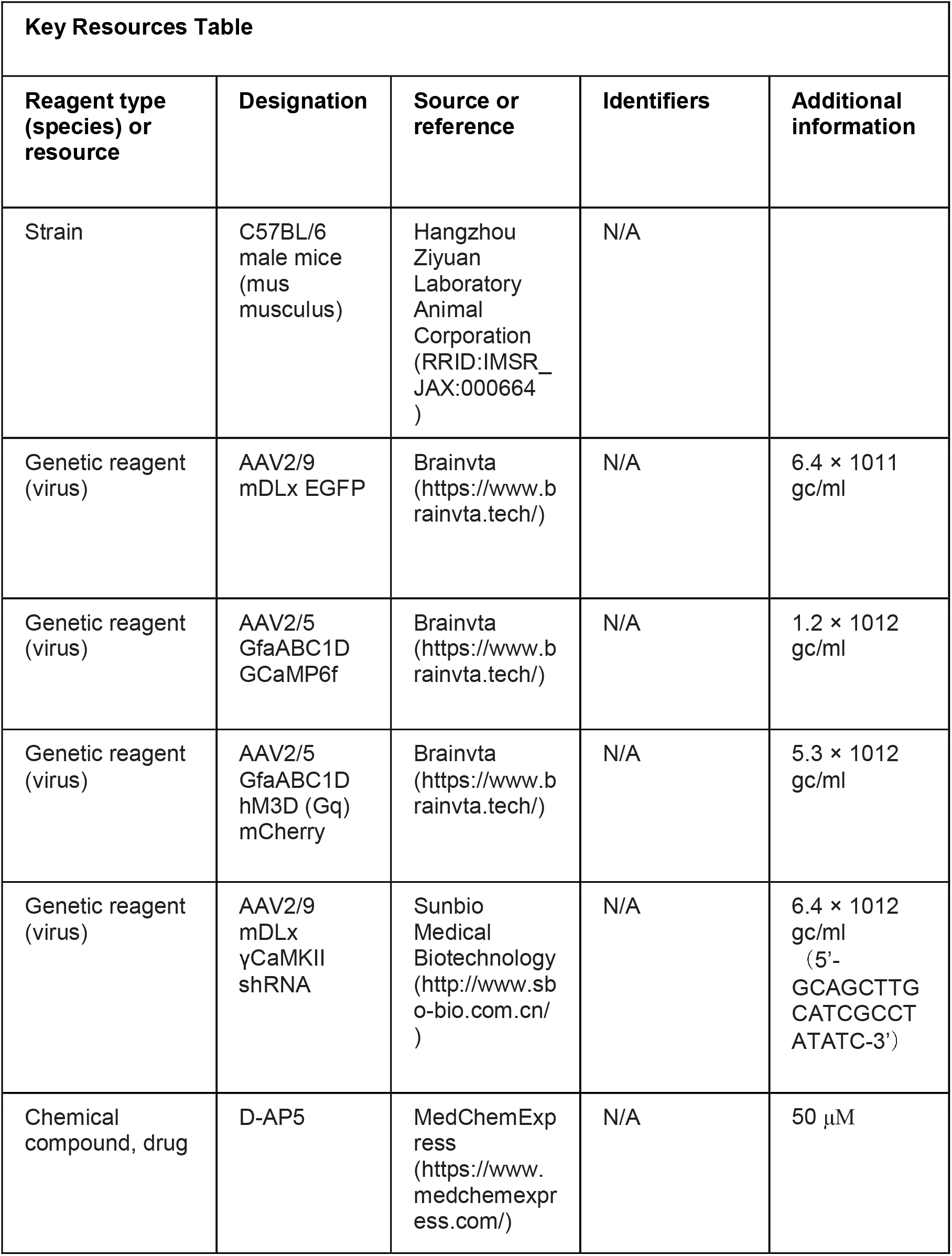
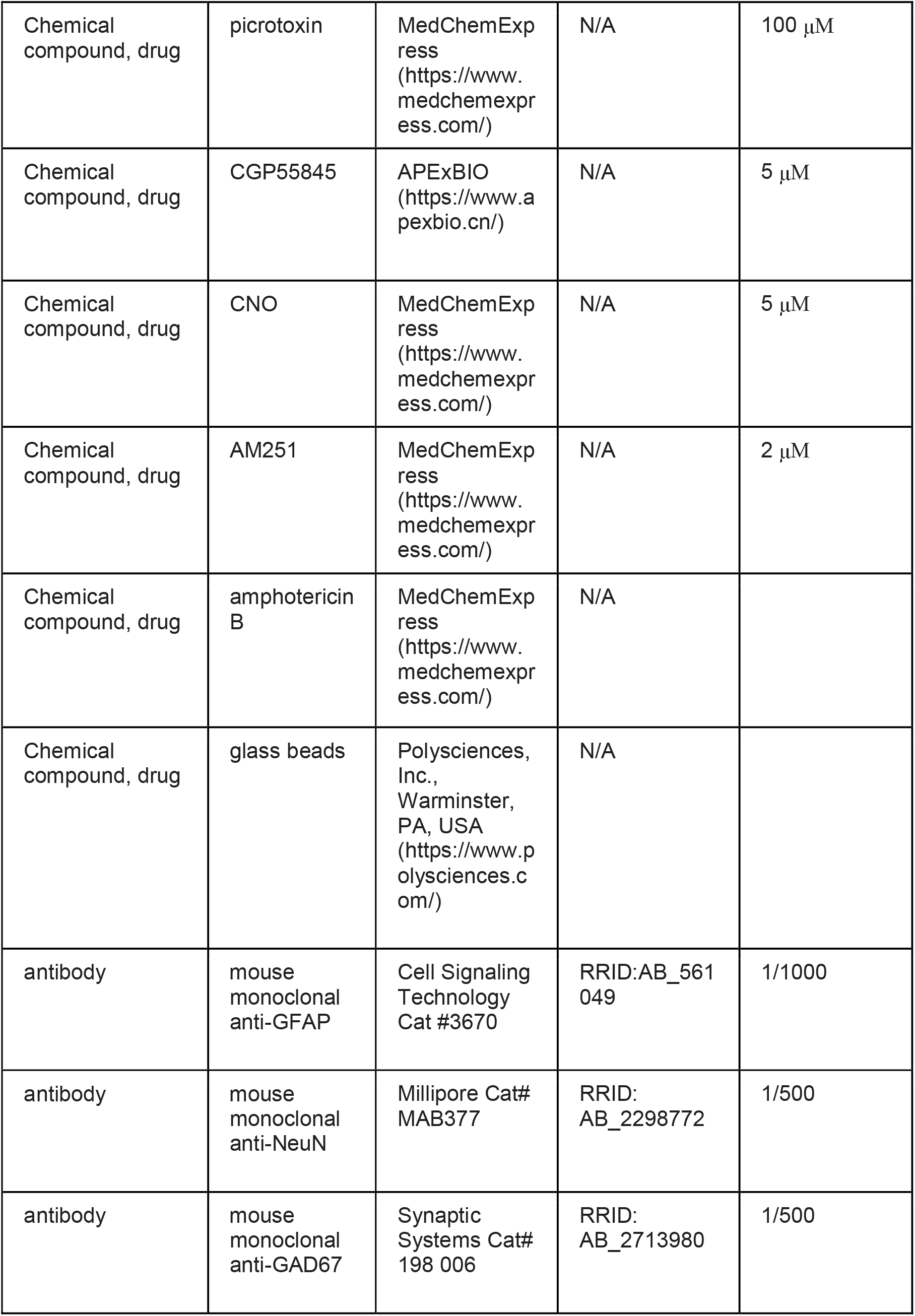
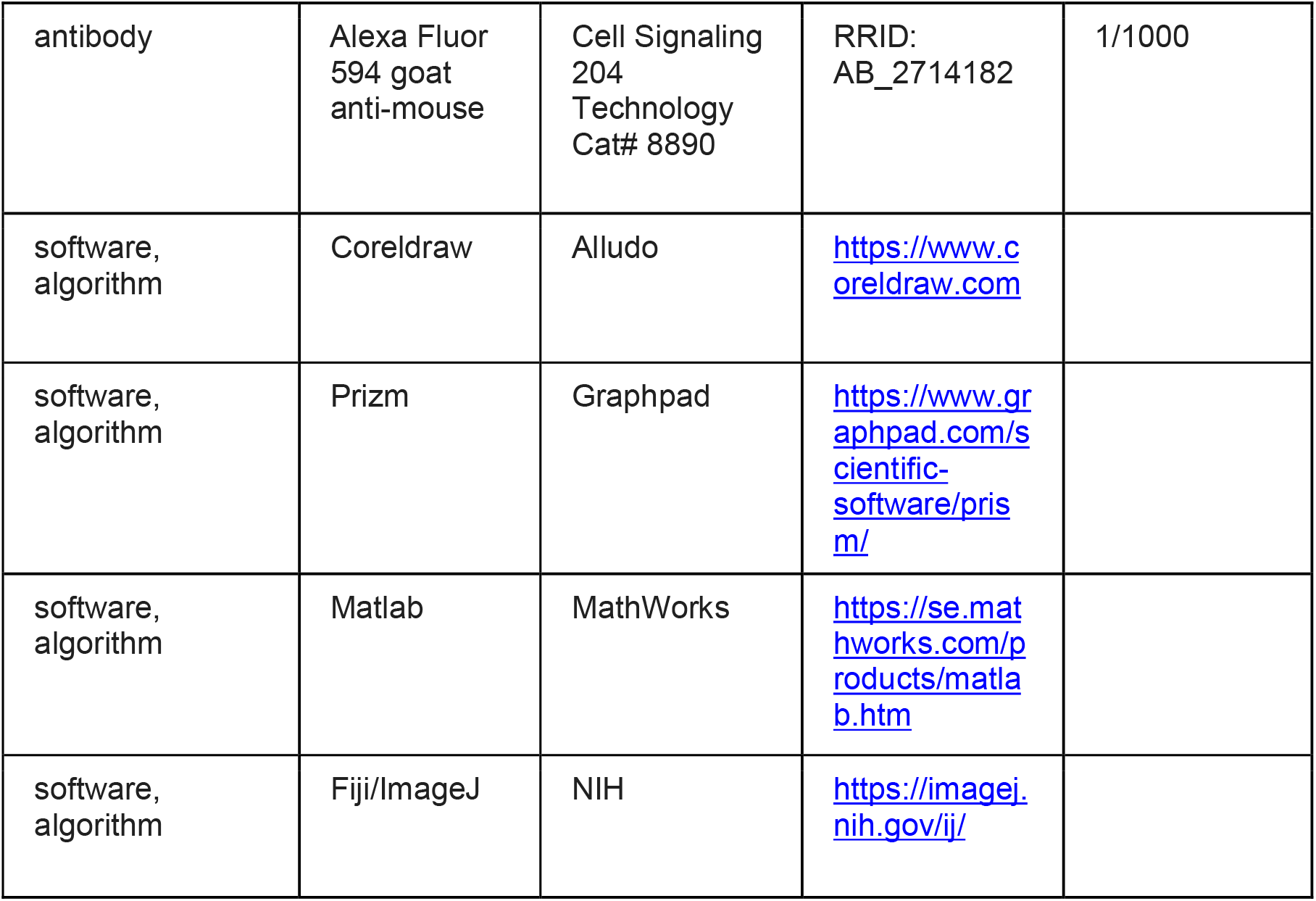

## Data availability statement

Source Data for all figures is available.

## Figure supplement

**Figure 1-figure supplement 1.**
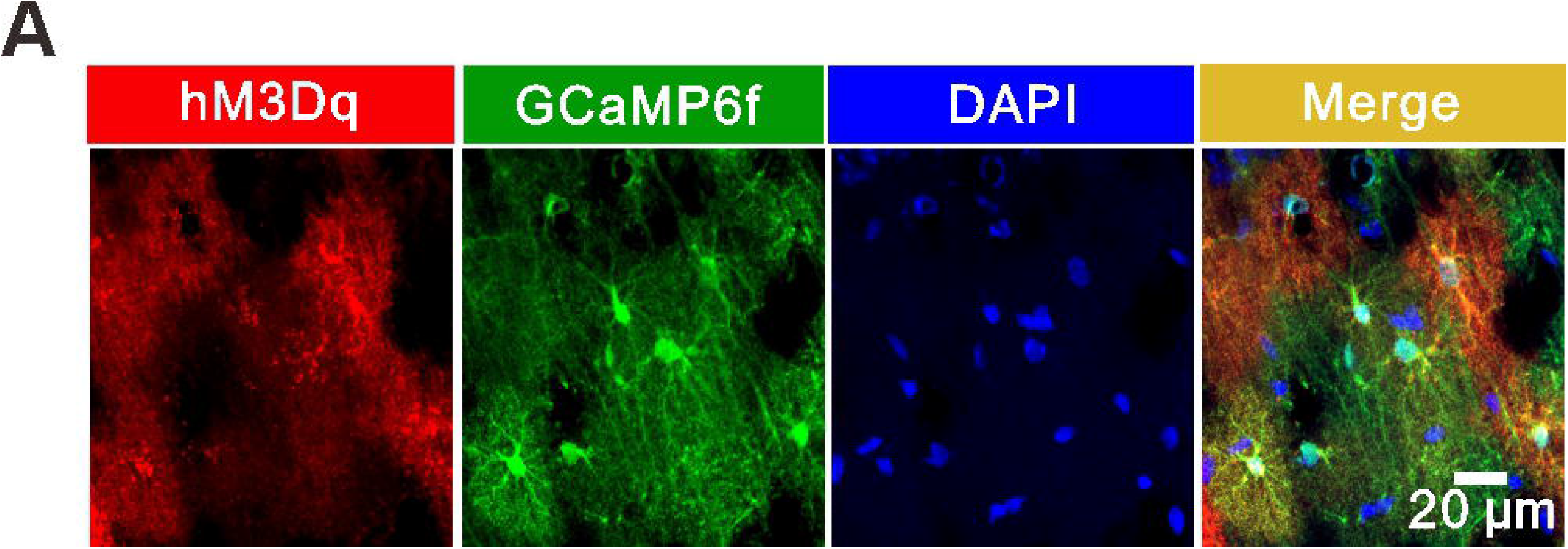
EGFP is expressed in GABAergic interneurons in the stratum radiatum of the hippocampus. (A) Schematic illustrating the procedure of AAV2/9 microinjections into the stratum radiatum of the hippocampus. (B) Confocal images showing EGFP (green) and GAD67 (red, white arrow)-expressing cells in the stratum radiatum of the hippocampus. (C and D) mDLx::EGFP was expressed in >98% of interneurons in the stratum radiatum of the hippocampus, with >98% specificity.

**Figure 1-figure supplement 2.**
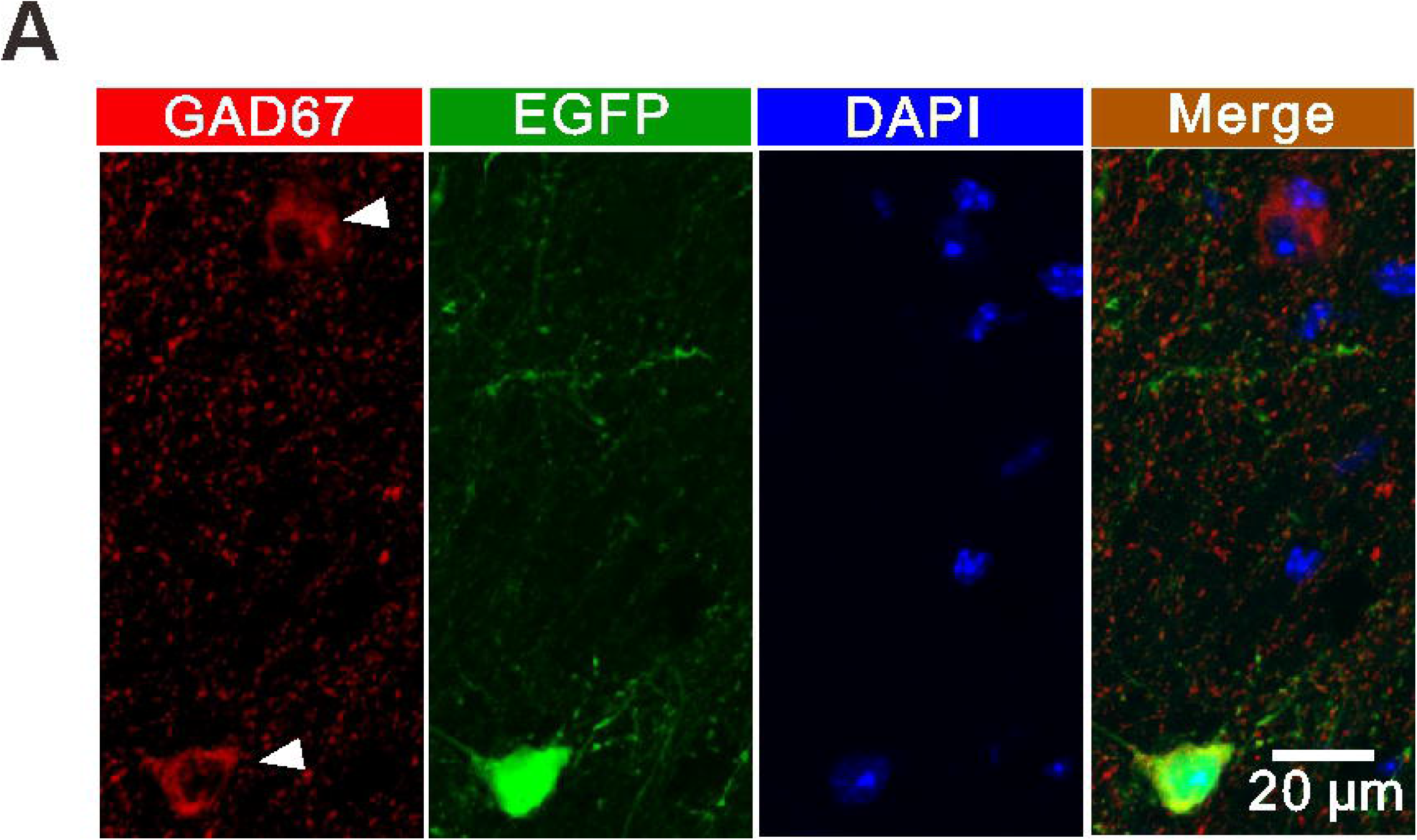
EGFP^+^ interneurons have normal sEPSCs, excitability and PPR. (A) Relationship of injected current to number of APs in postulated interneurons and EGFP^+^ interneurons. Inset: traces of membrane responses to current injection. (B) The number of APs at up state membrane potential (Vm) values **(Control: 14.62±1.474, n=13 slices from 6 mice; EGFP: 15.61±1.458, n=18 slices from 6 mice; t=0.4684 with 29 degrees of freedom, *p*=0.643, two-tailed unpaired t-test).** (C) Interneuron resting membrane potentials **(Control: -63.85±1.779 mV, n=13 slices from 6 mice; EGFP: -61.44±1.539 mV, n=18 slices from 6 mice; t=1.018 with 29 degrees of freedom, *p*=0.317, two-tailed unpaired t-test).** (D) Left: example traces of sEPSCs measured in the stratum radiatum of hippocampal postulated interneurons of WT mice; Right: example traces of sEPSCs measured in the stratum radiatum of hippocampal EGFP^+^ interneurons of virus-injected mice. (E, F) Cumulative distribution plots and summary of sEPSC amplitude and frequency in postulated interneurons and EGFP^+^ interneurons **(Amplitude: 16.89±1.846 pA of control, 18.28±2.003 of EGFP, n=10 slices from 6 mice, t=0.5125 with 18 degrees of freedom, *p*=0.6146, two-tailed unpaired t-test; Frequency: 12.41±3.058 Hz of control, 13.78±2.601 Hz of EGFP, n=10 slices from 6 mice, t=0.3405 with 18 degrees of freedom, *p*=0.7374, two-tailed unpaired t-test).** (G) Example paired-pulse traces measured in postulated interneurons and EGFP^+^ interneurons. (H) Summary of the paired-pulse ratio **(Control: 1.602±0.1443, n=8 slices from 3 mice; EGFP: 1.677±0.1794, n=8 slices from 3 mice; t=0.3267 with 14 degrees of freedom, *p*=0.7488, two-tailed unpaired t-test).**

**Figure 1-figure supplement 3.**
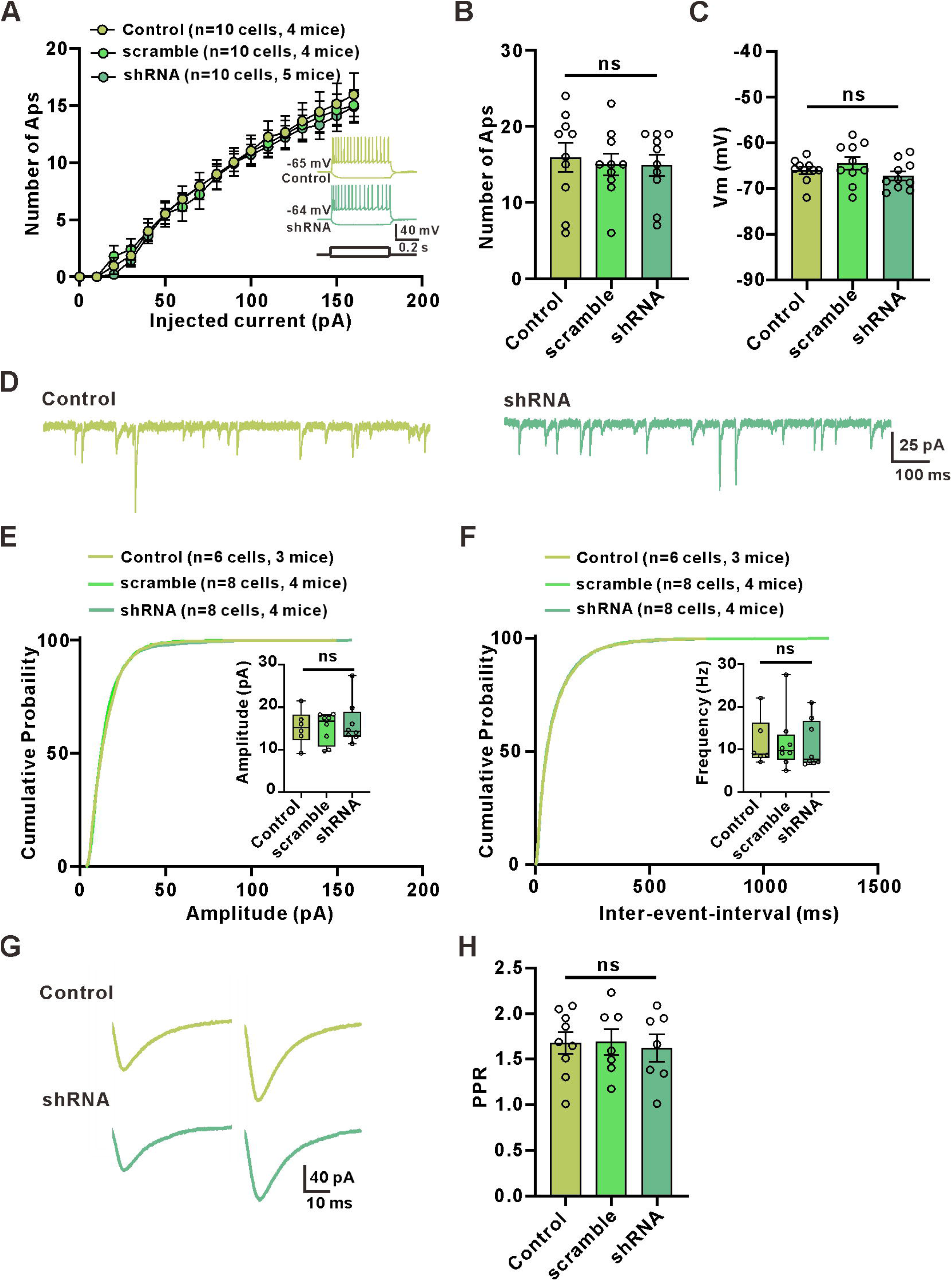
Stratum radiatum interneurons show linear rectifying AMPARs and a large NMDAR-mediated component. (A) Current-voltage (I-V) relation of AMPAR-mediated EPSCs in stratum radiatum interneurons. Inset: averaged EPSC traces at -90, -60, -30, 0 and 60 mV, showing the times at which the two components were measured. (B) The AMPA rectification index (EPSC amplitude at 60 mV / EPSC amplitude at - 60 mV) in eight interneurons from the stratum radiatum indicates linear rectifying AMPARs **(AMPA retification index: 0.8959±0.09362)**. (C) I-V relation for the NMDAR-mediated EPSCs in stratum radiatum interneurons. (D) NMDA/AMPA ratio (NMDAR-mediated EPSC amplitude at 60 mV / AMPAR-mediated EPSC amplitude at -60 mV) in eight cells from the stratum radiatum **(NMDA/AMPA ratio: 0.8664±0.2075)**.

**Figure 1-figure supplement 4.**
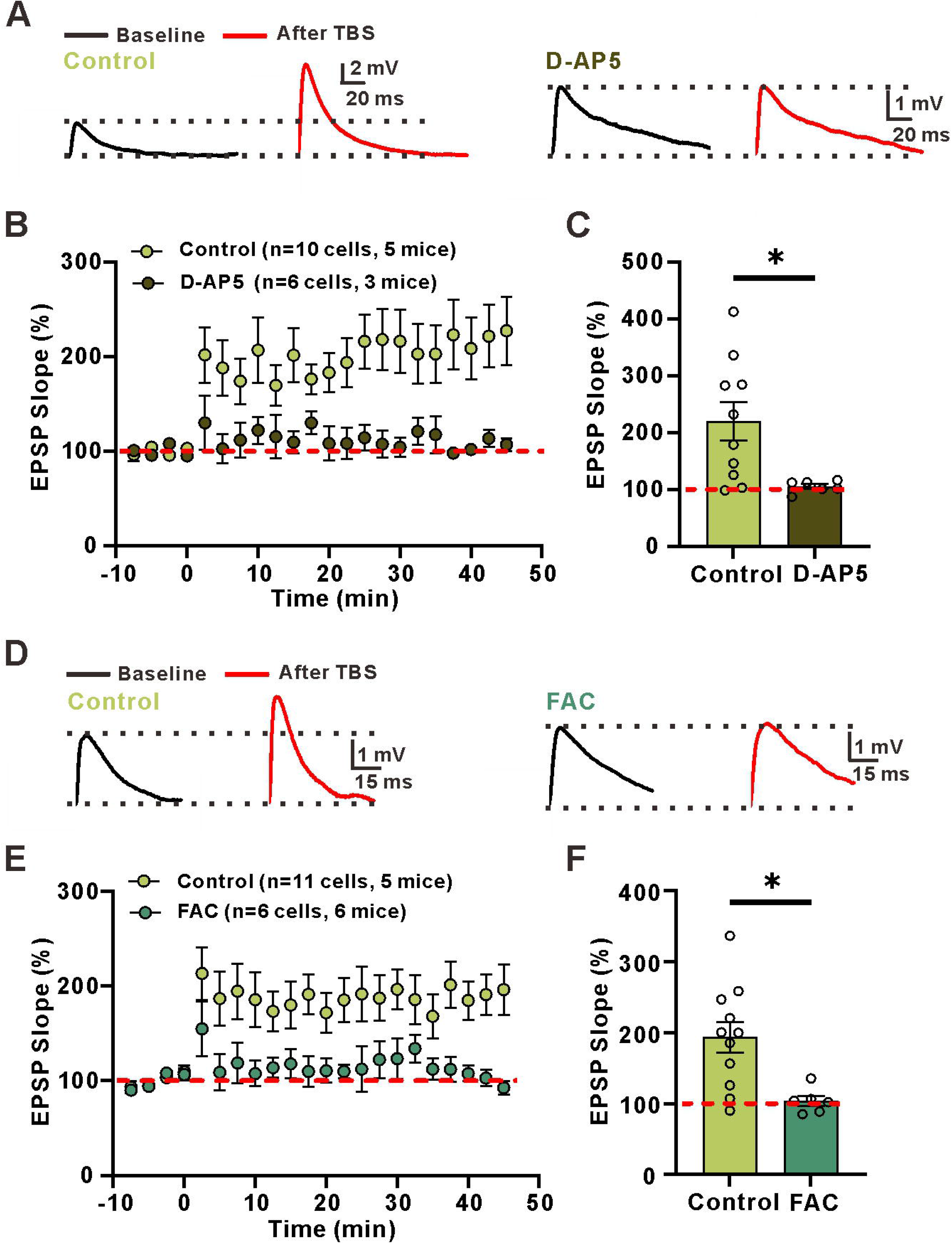
LTP_E→I_ in the stratum oriens is not dependent on astrocytic metabolism or the activation of NMDA receptors. (A) Left: superimposed representative averaged EPSPs recorded 10 min before (dark traces) and 35-45 min after (red traces) LTP induction; Right: superimposed representative averaged EPSPs recorded 10 min before (dark traces) and 35-45 min after (red traces) LTP induction when astrocytic metabolism was disrupted. (B) Normalized slope before and after the TBS stimulation protocol in control conditions and in the presence of the astrocytic metabolism inhibitor FAC. (C) The summary data measured 35-45 min after LTP induction **(Control: 233±52.74, n=5 slices from 3 mice, FAC: 233.5±33.06, n=6 slices from 6 mice; D-AP5:220.9±38.94, n=8 slices from 4 mice; *p*=0.96, F(2, 16)=0.03277, ANOVA with Dunnett’s comparison).**

**Figure 1-figure supplement 5.**
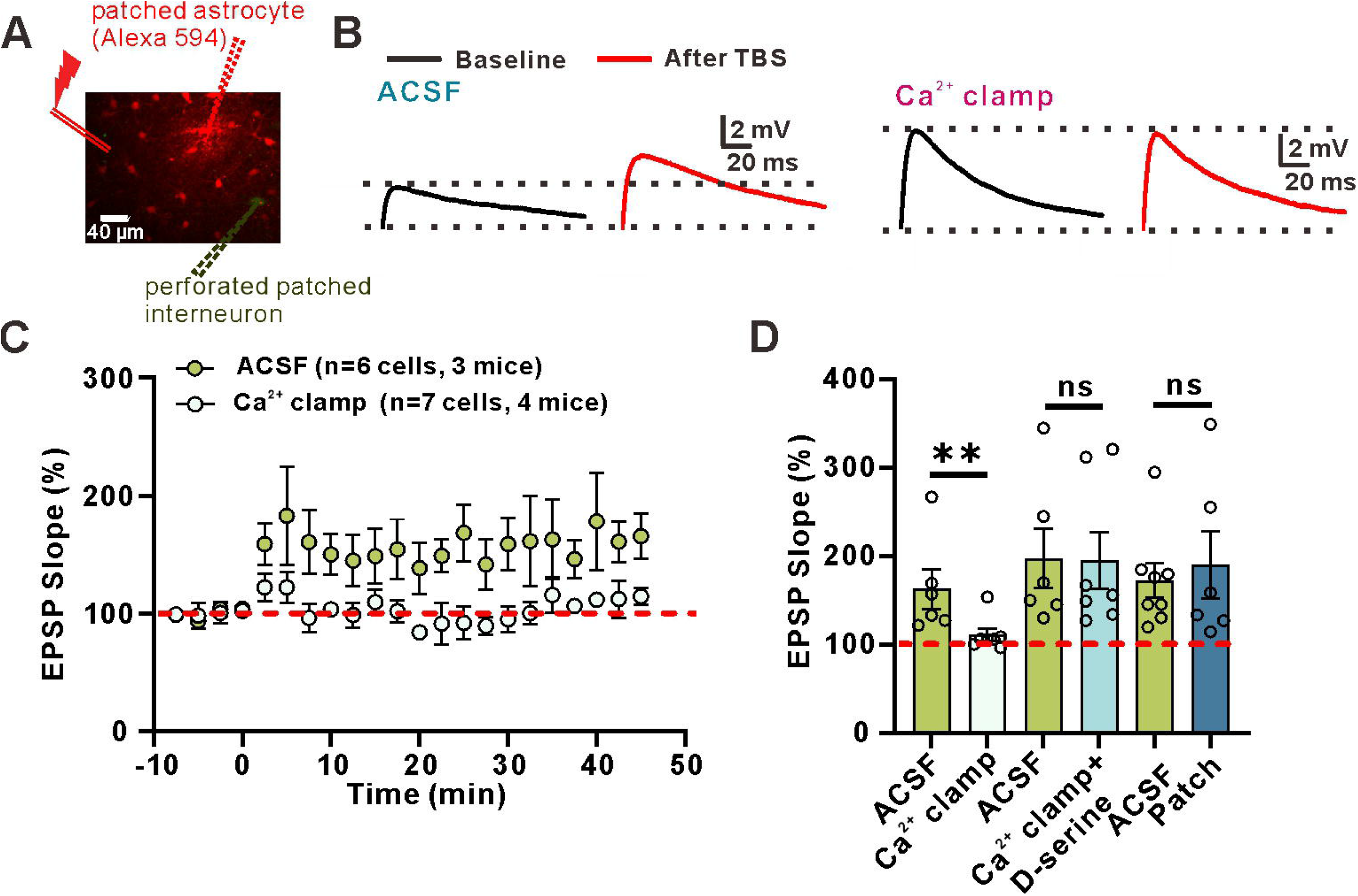
Stratum oriens interneurons show inwardly rectifying AMPARs and a negligible NMDAR-mediated component. (A) Current-voltage (I-V) relation of AMPAR-mediated EPSCs in stratum oriens interneurons. Inset: averaged EPSC traces at -90, -60, -30, 0 and 60 mV, showing the times at which the two components were measured. (B) The AMPA rectification index (EPSC amplitude at 60 mV / EPSC amplitude at - 60 mV) in six interneurons from the stratum oriens indicates inwardly rectifying AMPARs **(AMPA retification index: 0.2406±0.05594)**. (C) I-V relation for the NMDAR-mediated EPSCs in stratum oriens interneurons. (D) NMDA/AMPA ratio (NMDAR-mediated EPSC amplitude at 60 mV / AMPAR-mediated EPSC amplitude at -60 mV) in six cells from the stratum oriens **(NMDA/AMPA ratio: 0.1132±0.05370)**.

**Figure 3-figure supplement 1.**
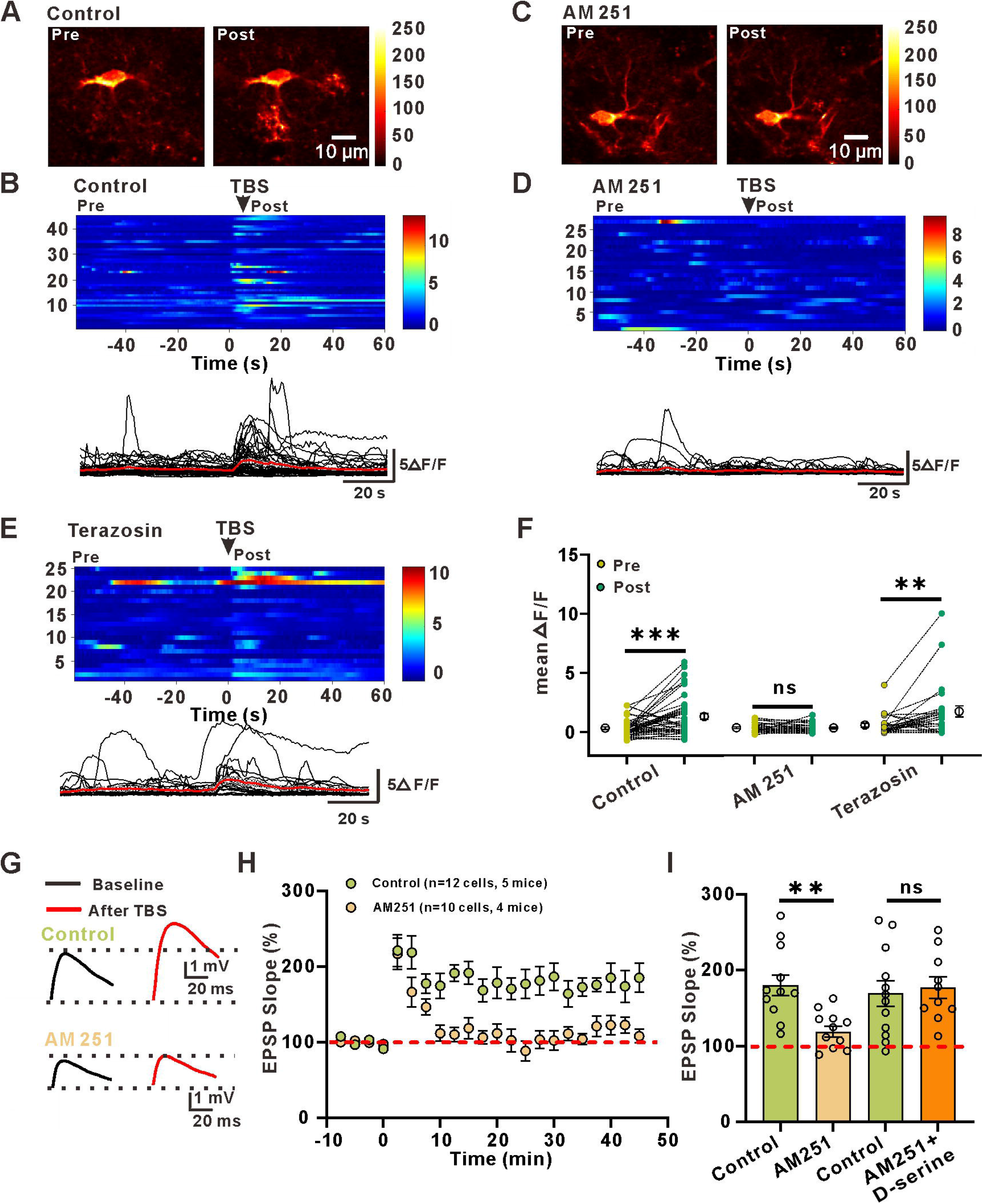
GCaMP6f was expressed in astrocytes. (A and B) Representative immunohistochemistry images showing colocalization of GCaMP6f and GFAP (A), but not NeuN (B). (C and D) gfaABC_1_D::GCaMP6f was expressed in >70% of astrocytes in the stratum radiatum of the hippocampus, with >99% specificity. (E) Quantification of the percentage of cells expressing NeuN that also expressed GCaMP6f.

**Figure 5-figure supplement 1.**
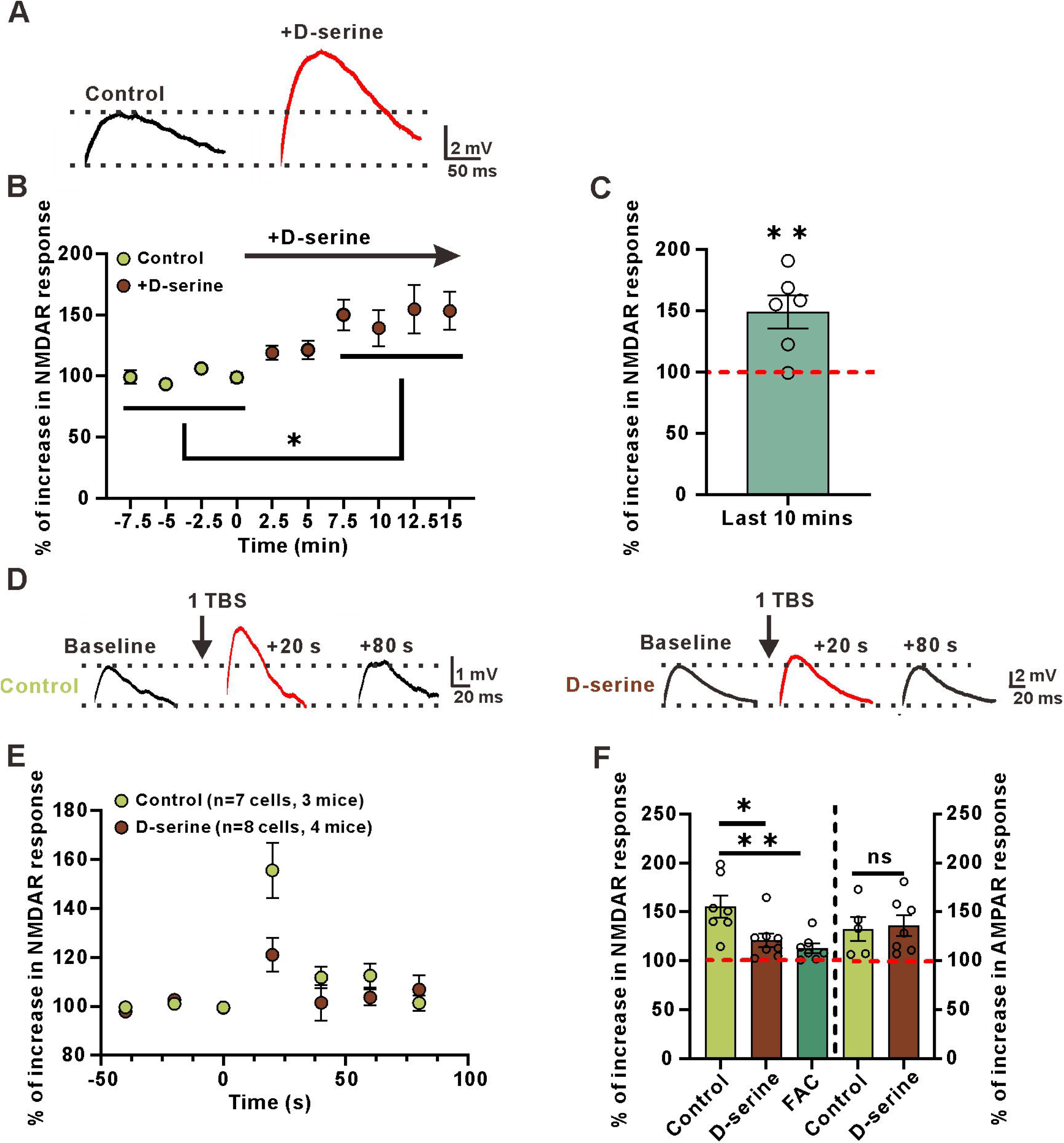
hM3Dq was expressed in astrocytes. (A) hM3Dq(mCherry) colocalization with GCaMP6f in the stratum radiatum of the hippocampus.

**Figure 6-figure supplement 1.**
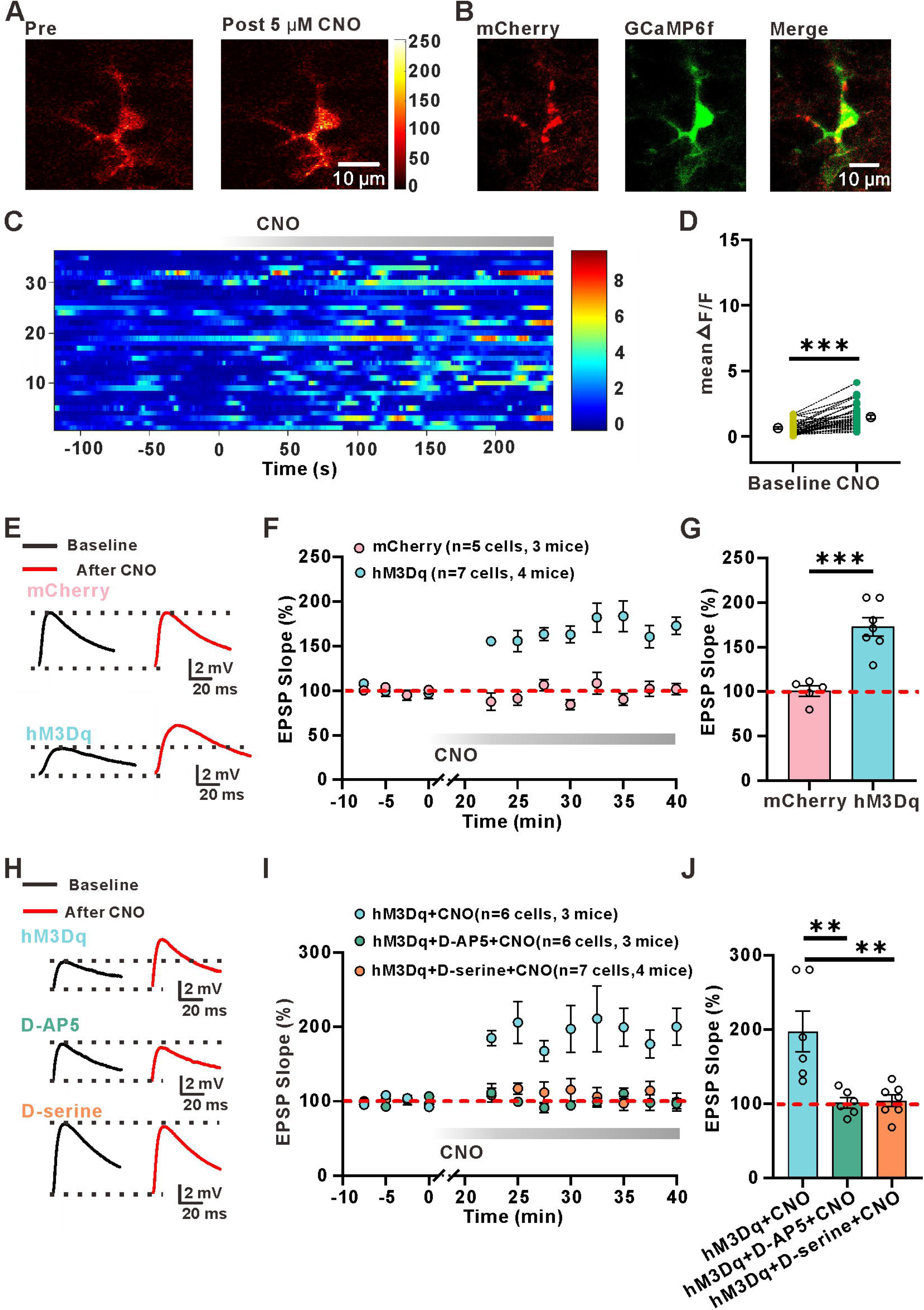
EGFP-γCaMKII shRNA is expressed in GABAergic interneurons in the stratum radiatum of the hippocampus. (A)Confocal images showing EGFP γCaMKII shRNA (green) and GAD67 (red, white arrows)-expressing cells in the stratum radiatum of the hippocampus. These images indicate that the γCaMKII shRNA, which is driven by an interneuron-specific promoter (mDLx), is expressed specifically in interneurons.

**Figure 6-figure supplement 2.**
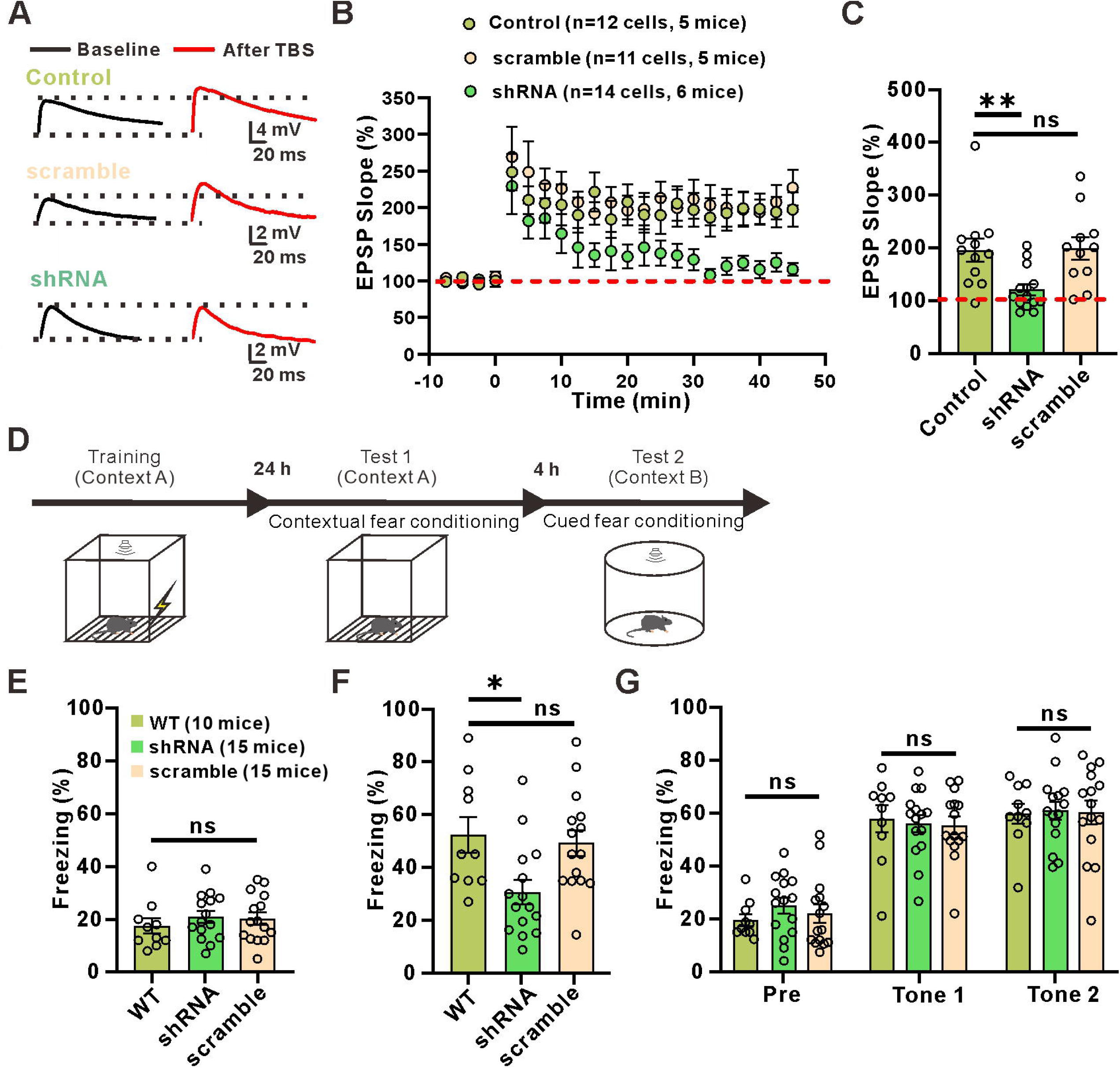
Knockdown of γCaMKII in interneurons has no effect on sEPSCs, excitability or the PPR. (A) Relationship of injected current to number of APs in postulated interneurons expressing scramble shRNA and γCaMKII shRNA interneurons. Inset: traces of membrane responses to current injection. (B) The number of APs at up state membrane potential (Vm) values **(Control: 15.95±1.904, n=10 slices from 4 mice; scramble shRNA: 15.00±1.430, n=10 slices from 4 mice;** γ**CaMKII shRNA: 14.90±1.410, n=10 slice from 5 mice; p=0.4986, F(2, 27)=0.4756, ANOVA with Dunnett’s comparison)**. (C) Interneuron resting membrane potentials **(Control: -66.03±0.8127, n=10 slices from 4 mice; scramble shRNA: -64.44±1.398, n=10 slices from 4 mice; γCaMKII shRNA: -67.23±0.9803, n=10 slice from 5 mice; p=0.2128, F(2, 27)=1.639, ANOVA with Dunnett’s comparison)**. (D) Left: example traces of sEPSCs measured in the stratum radiatum of hippocampal postulated interneurons of WT mice; Right: example traces of sEPSCs measured in the stratum radiatum of hippocampal EGFP^+^ interneurons of AAV-mDLx-shRNA γCaMKII-expressing mice. (E, F) Cumulative distribution plots and summary of sEPSC amplitude and frequency in postulated interneurons expressing scramble shRNA and γCaMKII shRNA interneurons **(Frequency: 11.62±2.328 Hz of control, n=6 slices from 3 mice, 11.69±2.462 Hz of scramble shRNA, n=8 slices from 4 mice, 11.12±2.015 Hz of** γ**CaMKII shRNA, n=8 slices from 4 mice, p=0.9805, F(2, 19)=0.01969, ANOVA with Dunnett’s comparison; Amplitude: 15.23±1.846 pA of control, n=6 slices from 3 mice, 14.57±1.382 of scramble shRNA, n=8 slices from 4 mice,16.23±1.821 of** γ**CaMKII shRNA, n=8 slices from 4 mice, p=0.8459, F(2, 19)=0.1688, ANOVA with Dunnett’s comparison)**. (G) Example paired-pulse traces measured in postulated interneurons of WT mice and EGFP^+^ interneurons of AAV-mDLx-shRNA γCaMKII-expressing mice. (H) Summary of the paired-pulse ratio **(Control: 1.679±0.1190 n=9 slices from 3 mice; scramble shRNA: 1.688±0.1405, n=7 slices from 3 mice; γCaMKII shRNA: 1.622±0.1477, n=7 slice from 3 mice; p=0.9361, F(2, 20)=0.06628, ANOVA with Dunnett’s comparison).**

Figure 1—source data 1

**LTP_E → I_ in the stratum radiatum is dependent on the activation of NMDA receptors and astrocytic metabolism.**

Figure 1—figure supplement 1—source data 1

**EGFP is expressed in GABAergic interneurons in the stratum radiatum of the hippocampus.**

Figure 1—figure supplement 2—source data 1

**EGFP^+^ interneurons have normal sEPSCs, excitability and PPR.**

Figure 1—figure supplement 3—source data 1

**Stratum radiatum interneurons show linear rectifying AMPARs and a large NMDAR-mediated component.**

Figure 1—figure supplement 4—source data 1

**LTP_E→I_ in the stratum oriens is not dependent on astrocytic metabolism or the activation of NMDA receptors.**

Figure 1—figure supplement 5—source data 1

**Stratum oriens interneurons show inwardly rectifying AMPARs and a negligible NMDAR-mediated component.**

Figure 2—source data 1

**Astrocyte Ca^2+^ is involved in the induction of LTP.**

Figure 3—source data 1

**Activation of astrocytic CB1 receptors causes an increase in astrocytic Ca^2+^ signals and is involved in the induction of LTP_E→I_.**

Figure 3—figure supplement 1—source data 1

**GCaMP6f was expressed in astrocytes.**

Figure 4—source data 1

**D-Serine release from astrocytes potentiates NMDAR-mediated responses.**

Figure 5—source data 1

**Astrocytic activation induces de novo LTP_E_**_→_**_I_.**

Figure 5—figure supplement 1—source data 1

**hM3Dq was expressed in astrocytes.**

Figure 6—source data 1

**Impaired hippocampus-dependent long-term memory in γCaMKII knockdown mice.**

Figure 6—figure supplement 1—source data 1

**Knockdown of γCaMKII in interneurons has no effect on sEPSCs, excitability or the PPR.**

## References

Adamsky, A., Kol, A., Kreisel, T. et al. (2018) Astrocytic Activation Generates De Novo Neuronal Potentiation and Memory Enhancement. Cell 174, 59–71 e14.

Araque, A., Carmignoto, G., Haydon, P. G., Oliet, S. H., Robitaille, R. and Volterra, A. (2014) Gliotransmitters travel in time and space. Vol. 81, pp. 728–739. Neuron.

Asgarihafshejani, A., Honore, E., Michon, F. X., Laplante, I. and Lacaille, J. C. (2022) Long-term potentiation at pyramidal cell to somatostatin interneuron synapses controls hippocampal network plasticity and memory. iScience 25, 104259.

Bacci, A., Huguenard, J. R. and Prince, D. A. (2004) Long-lasting self-inhibition of neocortical interneurons mediated by endocannabinoids. Nature 431, 312–316.

Bazargani, N. and Attwell, D. (2016) Astrocyte calcium signaling: the third wave. Nat. Neurosci. 19, 182–189.

Bekar, L. K., He, W. and Nedergaard, M. (2008) Locus coeruleus alpha-adrenergic-mediated activation of cortical astrocytes in vivo. Cerebral cortex (New York, N.Y. : 1991) 18, 2789–2795.

Benneyworth, M. A., Li, Y., Basu, A. C., Bolshakov, V. Y. and Coyle, J. T. (2012) Cell selective conditional null mutations of serine racemase demonstrate a predominate localization in cortical glutamatergic neurons. Cell. Mol. Neurobiol. 32, 613–624.

Bezaire, M. J. and Soltesz, I. (2013) Quantitative assessment of CA1 local circuits: knowledge base for interneuron-pyramidal cell connectivity. Hippocampus 23, 751–785.

Bliss, T. V. and Collingridge, G. L. (1993) A synaptic model of memory: long-term potentiation in the hippocampus. Nature 361, 31–39.

Bliss, T. V. and Lomo, T. (1973) Long-lasting potentiation of synaptic transmission in the dentate area of the anaesthetized rabbit following stimulation of the perforant path. J. Physiol. 232, 331–356.

Bohmbach, K., Masala, N., Schonhense, E. M., Hill, K., Haubrich, A. N., Zimmer, A., Opitz, T., Beck, H. and Henneberger, C. (2022) An astrocytic signaling loop for frequency-dependent control of dendritic integration and spatial learning. Nat Commun 13, 7932.

Booker, S. A. and Vida, I. (2018) Morphological diversity and connectivity of hippocampal interneurons. Cell Tissue Res. 373, 619–641.

Bushong, E. A., Martone, M. E., Jones, Y. Z. and Ellisman, M. H. (2002) Protoplasmic astrocytes in CA1 stratum radiatum occupy separate anatomical domains. J. Neurosci. 22, 183–192.

Castillo, P. E., Younts, T. J., Chavez, A. E. and Hashimotodani, Y. (2012) Endocannabinoid signaling and synaptic function. Neuron 76, 70–81.

Chaboub, L. S. and Deneen, B. (2013) Astrocyte form and function in the developing central nervous system. Semin. Pediatr. Neurol. 20, 230–235.

Ciappelloni, S., Murphy-Royal, C., Dupuis, J. P., Oliet, S. H. R. and Groc, L. (2017) Dynamics of surface neurotransmitter receptors and transporters in glial cells: Single molecule insights. Cell Calcium 67, 46–52.

Covelo, A. and Araque, A. (2018) Neuronal activity determines distinct gliotransmitter release from a single astrocyte. Elife 7.

De Pitta, M., Brunel, N. and Volterra, A. (2016) Astrocytes: Orchestrating synaptic plasticity? Neuroscience 323, 43–61.

Dimidschstein, J., Chen, Q., Tremblay, R. et al. (2016) A viral strategy for targeting and manipulating interneurons across vertebrate species. Nat. Neurosci. 19, 1743–1749.

Ding, F., O’Donnell, J., Thrane, A. S., Zeppenfeld, D., Kang, H., Xie, L., Wang, F. and Nedergaard, M. (2013) alpha1-Adrenergic receptors mediate coordinated Ca2+ signaling of cortical astrocytes in awake, behaving mice. Cell Calcium 54, 387–394.

Dun, Y., Duplantier, J., Roon, P., Martin, P. M., Ganapathy, V. and Smith, S. B. (2008) Serine racemase expression and D-serine content are developmentally regulated in neuronal ganglion cells of the retina. J. Neurochem. 104, 970–978.

Eraso-Pichot, A., Pouvreau, S., Olivera-Pinto, A., Gomez-Sotres, P., Skupio, U. and Marsicano, G. (2023) Endocannabinoid signaling in astrocytes. Glia 71, 44–59.

Escartin, C., Guillemaud, O. and Carrillo-de Sauvage, M. A. (2019) Questions and (some) answers on reactive astrocytes. Glia 67, 2221–2247.

Fernandez-Moncada, I. and Marsicano, G. (2023) Astroglial CB1 receptors, energy metabolism, and gliotransmission: an integrated signaling system? Essays Biochem. 67, 49–61.

Goenaga, J., Araque, A., Kofuji, P. and Herrera Moro Chao, D. (2023) Calcium signaling in astrocytes and gliotransmitter release. Front. Synaptic Neurosci. 15, 1138577.

Gordon, G. R., Baimoukhametova, D. V., Hewitt, S. A., Rajapaksha, W. R., Fisher, T. E. and Bains, J. S. (2005) Norepinephrine triggers release of glial ATP to increase postsynaptic efficacy. Nat. Neurosci. 8, 1078–1086.

Gutierrez-Rodriguez, A., Bonilla-Del Rio, I., Puente, N. et al. (2018) Localization of the cannabinoid type-1 receptor in subcellular astrocyte compartments of mutant mouse hippocampus. Glia 66, 1417–1431.

Hamilton, N. B. and Attwell, D. (2010) Do astrocytes really exocytose neurotransmitters? Nat. Rev. Neurosci. 11, 227–238.

Han, J., Kesner, P., Metna-Laurent, M. et al. (2012) Acute cannabinoids impair working memory through astroglial CB1 receptor modulation of hippocampal LTD. Cell 148, 1039–1050.

He, X., Li, J., Zhou, G. et al. (2021) Gating of hippocampal rhythms and memory by synaptic plasticity in inhibitory interneurons. Neuron 109, 1013–1028 e1019.

He, X., Wang, Y., Zhou, G., Yang, J., Li, J., Li, T., Hu, H. and Ma, H. (2022) A Critical Role for gammaCaMKII in Decoding NMDA Signaling to Regulate AMPA Receptors in Putative Inhibitory Interneurons. Neurosci. Bull. 38, 916–926.

Henneberger, C., Papouin, T., Oliet, S. H. and Rusakov, D. A. (2010) Long-term potentiation depends on release of D-serine from astrocytes. Nature 463, 232–236.

Hosli, L., Binini, N., Ferrari, K. D. et al. (2022) Decoupling astrocytes in adult mice impairs synaptic plasticity and spatial learning. Cell Rep. 38, 110484.

Jimenez-Blasco, D., Busquets-Garcia, A., Hebert-Chatelain, E. et al. (2020) Glucose metabolism links astroglial mitochondria to cannabinoid effects. Nature 583, 603–608.

Kano, M., Ohno-Shosaku, T., Hashimotodani, Y., Uchigashima, M. and Watanabe, M. (2009) Endocannabinoid-mediated control of synaptic transmission. Physiol. Rev. 89, 309–380.

Khakh, B. S. and McCarthy, K. D. (2015) Astrocyte calcium signaling: from observations to functions and the challenges therein. Cold Spring Harb. Perspect. Biol. 7, a020404.

Kullmann, D. M. and Lamsa, K. P. (2007) Long-term synaptic plasticity in hippocampal interneurons. Nat. Rev. Neurosci. 8, 687–699.

Kullmann, D. M. and Lamsa, K. P. (2011) LTP and LTD in cortical GABAergic interneurons: emerging rules and roles. Neuropharmacology 60, 712–719.

Lamsa, K., Heeroma, J. H. and Kullmann, D. M. (2005) Hebbian LTP in feed-forward inhibitory interneurons and the temporal fidelity of input discrimination. Nat. Neurosci. 8, 916–924.

Lamsa, K. P., Heeroma, J. H., Somogyi, P., Rusakov, D. A. and Kullmann, D. M. (2007) Anti-Hebbian long-term potentiation in the hippocampal feedback inhibitory circuit. Science 315, 1262–1266.

Le Duigou, C., Savary, E., Kullmann, D. M. and Miles, R. (2015) Induction of Anti-Hebbian LTP in CA1 Stratum Oriens Interneurons: Interactions between Group I Metabotropic Glutamate Receptors and M1 Muscarinic Receptors. J. Neurosci. 35, 13542–13554.

Liu, Y. Z., Wang, Y., Shen, W. and Wang, Z. (2017) Enhancement of synchronized activity between hippocampal CA1 neurons during initial storage of associative fear memory. J. Physiol. 595, 5327–5340.

Ma, W., Si, T., Wang, Z., Wang, J., Xu, F. and Li, Q. (2022) Astrocytic α4nAchRs Signaling in the Hippocampus Governs the Formation of Temporal Association Memory. SSRN Electronic Journal.

Marinelli, S., Pacioni, S., Cannich, A., Marsicano, G. and Bacci, A. (2009) Self-modulation of neocortical pyramidal neurons by endocannabinoids. Nat. Neurosci. 12, 1488–1490.

Miya, K., Inoue, R., Takata, Y., Abe, M., Natsume, R., Sakimura, K., Hongou, K., Miyawaki, T. and Mori, H. (2008) Serine racemase is predominantly localized in neurons in mouse brain. J. Comp. Neurol. 510, 641–654.

Mothet, J. P., Rouaud, E., Sinet, P. M., Potier, B., Jouvenceau, A., Dutar, P., Videau, C., Epelbaum, J. and Billard, J. M. (2006) A critical role for the glial-derived neuromodulator D-serine in the age-related deficits of cellular mechanisms of learning and memory. Aging Cell 5, 267–274.

Nam, M. H., Han, K. S., Lee, J. et al. (2019) Activation of Astrocytic mu-Opioid Receptor Causes Conditioned Place Preference. Cell Rep. 28, 1154–1166 e1155.

Navarrete, M. and Araque, A. (2008) Endocannabinoids mediate neuron-astrocyte communication. Neuron 57, 883–893.

Navarrete, M. and Araque, A. (2010) Endocannabinoids potentiate synaptic transmission through stimulation of astrocytes. Neuron 68, 113–126.

Navarrete, M., Diez, A. and Araque, A. (2014) Astrocytes in endocannabinoid signalling. Philos. Trans. R. Soc. Lond. B Biol. Sci. 369, 20130599.

Navarrete, M., Perea, G., Fernandez de Sevilla, D., Gomez-Gonzalo, M., Nunez, A., Martin, E. D. and Araque, A. (2012) Astrocytes mediate in vivo cholinergic-induced synaptic plasticity. PLoS Biol. 10, e1001259.

Nicoll, R. A. (2017) A Brief History of Long-Term Potentiation. Neuron 93, 281–290.

Nikolic, L., Shen, W., Nobili, P., Virenque, A., Ulmann, L. and Audinat, E. (2018) Blocking TNFalpha-driven astrocyte purinergic signaling restores normal synaptic activity during epileptogenesis. Glia 66, 2673–2683.

Nissen, W., Szabo, A., Somogyi, J., Somogyi, P. and Lamsa, K. P. (2010) Cell type-specific long-term plasticity at glutamatergic synapses onto hippocampal interneurons expressing either parvalbumin or CB1 cannabinoid receptor. J. Neurosci. 30, 1337–1347.

Noriega-Prieto, J. A., Kofuji, P. and Araque, A. (2023) Endocannabinoid signaling in synaptic function. Glia 71, 36–43.

Oe, Y., Wang, X., Patriarchi, T. et al. (2020) Distinct temporal integration of noradrenaline signaling by astrocytic second messengers during vigilance. Nat Commun 11, 471.

Oren, I., Nissen, W., Kullmann, D. M., Somogyi, P. and Lamsa, K. P. (2009) Role of ionotropic glutamate receptors in long-term potentiation in rat hippocampal CA1 oriens-lacunosum moleculare interneurons. J. Neurosci. 29, 939–950.

Panatier, A., Theodosis, D. T., Mothet, J. P., Touquet, B., Pollegioni, L., Poulain, D. A. and Oliet, S. H. (2006) Glia-derived D-serine controls NMDA receptor activity and synaptic memory. Cell 125, 775–784.

Pankratov, Y. and Lalo, U. (2015) Role for astroglial alpha1-adrenoreceptors in gliotransmission and control of synaptic plasticity in the neocortex. Front. Cell. Neurosci. 9, 230.

Papouin, T., Dunphy, J. M., Tolman, M., Dineley, K. T. and Haydon, P. G. (2017a) Septal Cholinergic Neuromodulation Tunes the Astrocyte-Dependent Gating of Hippocampal NMDA Receptors to Wakefulness. Neuron 94, 840–854 e847.

Papouin, T., Henneberger, C., Rusakov, D. A. and Oliet, S. H. R. (2017b) Astroglial versus Neuronal D-Serine: Fact Checking. Trends Neurosci. 40, 517–520.

Papouin, T., Ladepeche, L., Ruel, J. et al. (2012) Synaptic and extrasynaptic NMDA receptors are gated by different endogenous coagonists. Cell 150, 633–646.

Paukert, M., Agarwal, A., Cha, J., Doze, V. A., Kang, J. U. and Bergles, D. E. (2014) Norepinephrine controls astroglial responsiveness to local circuit activity. Neuron 82, 1263–1270.

Pelkey, K. A., Chittajallu, R., Craig, M. T., Tricoire, L., Wester, J. C. and McBain, C. J. (2017) Hippocampal GABAergic Inhibitory Interneurons. Physiol. Rev. 97, 1619–1747.

Pelletier, J. G. and Lacaille, J. C. (2008) Long-term synaptic plasticity in hippocampal feedback inhibitory networks. Prog. Brain Res. 169, 241–250.

Perez-Catalan, N. A., Doe, C. Q. and Ackerman, S. D. (2021) The role of astrocyte-mediated plasticity in neural circuit development and function. Neural Dev 16, 1.

Ramon-Duaso, C., Conde-Moro, A. R. and Busquets-Garcia, A. (2023) Astroglial cannabinoid signaling and behavior. Glia 71, 60–70.

Robin, L. M., Oliveira da Cruz, J. F., Langlais, V. C., et al. (2018) Astroglial CB(1) Receptors Determine Synaptic D-Serine Availability to Enable Recognition Memory. Neuron 98, 935–944 e935.

Sancho, L., Contreras, M. and Allen, N. J. (2021) Glia as sculptors of synaptic plasticity. Neurosci. Res. 167, 17–29.

Santello, M., Toni, N. and Volterra, A. (2019) Astrocyte function from information processing to cognition and cognitive impairment. Nat. Neurosci. 22, 154–166.

Shen, W., Chen, S., Xiang, Y., Yao, Z., Chen, Z., Wu, X., Li, L. and Zeng, L. H. (2021) Astroglial adrenoreceptors modulate synaptic transmission and contextual fear memory formation in dentate gyrus. Neurochem. Int. 143, 104942.

Shen, W., Li, Z., Tang, Y. et al. (2022) Somatostatin interneurons inhibit excitatory transmission mediated by astrocytic GABAB and presynaptic GABAB and adenosine A1 receptors in the hippocampus. J. Neurochem.

Shen, W., Nikolic, L., Meunier, C., Pfrieger, F. and Audinat, E. (2017) An autocrine purinergic signaling controls astrocyte-induced neuronal excitation. Sci. Rep. 7, 11280.

Sherwood, M. W., Arizono, M., Hisatsune, C., Bannai, H., Ebisui, E., Sherwood, J. L., Panatier, A., Oliet, S. H. and Mikoshiba, K. (2017) Astrocytic IP(3) Rs: Contribution to Ca(2+) signalling and hippocampal LTP. Glia 65, 502–513.

Stevens, E. R., Esguerra, M., Kim, P. M., Newman, E. A., Snyder, S. H., Zahs, K. R. and Miller, R. F. (2003) D-serine and serine racemase are present in the vertebrate retina and contribute to the physiological activation of NMDA receptors. Proc. Natl. Acad. Sci. U. S. A. 100, 6789–6794.

Swanson, R. A. and Graham, S. H. (1994) Fluorocitrate and fluoroacetate effects on astrocyte metabolism in vitro. Brain Res. 664, 94–100.

Van Den Herrewegen, Y., Sanderson, T. M., Sahu, S., De Bundel, D., Bortolotto, Z. A. and Smolders, I. (2021) Side-by-side comparison of the effects of Gq- and Gi-DREADD-mediated astrocyte modulation on intracellular calcium dynamics and synaptic plasticity in the hippocampal CA1. Mol. Brain 14, 144.

Verkhratsky, A. and Nedergaard, M. (2018) Physiology of Astroglia. Physiol. Rev. 98, 239–389.

Verkhratsky, A. and Steinhäuser, C. (2000) Ion channels in glial cells. Brain Res. Rev. 32, 380–412.

Wolosker, H., Balu, D. T. and Coyle, J. T. (2016) The Rise and Fall of the d-Serine-Mediated Gliotransmission Hypothesis. Trends Neurosci. 39, 712–721.

Wolosker, H., Balu, D. T. and Coyle, J. T. (2017) Astroglial Versus Neuronal D-Serine: Check Your Controls! Trends Neurosci. 40, 520–522.

Wolosker, H., Blackshaw, S. and Snyder, S. H. (1999) Serine racemase: a glial enzyme synthesizing D-serine to regulate glutamate-N-methyl-D-aspartate neurotransmission. Proc. Natl. Acad. Sci. U. S. A. 96, 13409–13414.

Xie, Y., Kuan, A. T., Wang, W. et al. (2022) Astrocyte-neuron crosstalk through Hedgehog signaling mediates cortical synapse development. Cell Rep. 38, 110416.

Yang, Y., Ge, W., Chen, Y., Zhang, Z., Shen, W., Wu, C., Poo, M. and Duan, S. (2003) Contribution of astrocytes to hippocampal long-term potentiation through release of D-serine. Proc. Natl. Acad. Sci. U. S. A. 100, 15194–15199.

